# Impact of high-fat diet on lifespan, metabolism, fecundity and behavioral senescence in *Drosophila*

**DOI:** 10.1101/2020.06.28.175794

**Authors:** Sifang Liao, Mirjam Amcoff, Dick R. Nässel

## Abstract

Excess consumption of high-fat diet (HFD) is likely to result in obesity and increases the predisposition to associated health disorders. *Drosophila melanogaster* has emerged as an important model to study the effects of HFD on metabolism, gut function, behavior, and ageing. In this study, we investigated the effects of HFD on physiology and behavior of female flies at different time-points over several weeks. We found that HFD decreases lifespan, and also with age leads to accelerated decline of climbing ability in both virgins and mated flies. In virgins HFD also increased sleep fragmentation with age. Furthermore, long-term exposure to HFD results in elevated adipokinetic hormone (AKH) transcript levels and an enlarged crop with increased lipid stores. We detected no long-term effects of HFD on body mass, or levels of triacylglycerides (TAG), glycogen or glucose, although fecundity was diminished. However, one week of HFD resulted in decreased body mass and elevated TAG levels in mated flies. Finally, we investigated the role of AKH in regulating effects of HFD during aging. Both with normal diet (ND) and HFD, *Akh* mutant flies displayed increased longevity compared to control flies. However, both mutants and controls showed shortened lifespan on HFD compared to ND. In flies exposed to ND, fecundity is decreased in *Akh* mutants compared to controls after one week, but increased after three weeks. However, HFD leads to a similar decrease in fecundity in both genotypes after both exposure times. Thus, long-term exposure to HFD increases AKH signaling, impairs lifespan and fecundity and augments age-related behavioral senescence.

## 1. Introduction

Intake of high-fat diet (HFD) is likely to result in obesity, which in turn increases the predisposition to cardiovascular disease, diabetes, cancer, and other metabolic disorders (Bray and Popkin, 1998; Dietrich et al., 2013; Hill et al., 2000; Musselman and Kühnlein, 2018; O’Brien and Dixon, 2002; Szendroedi and Roden, 2009; van Herpen and Schrauwen-Hinderling, 2008). Obesity is also linked to increased risk of cognitive impairment (Hwang et al., 2010; Liu et al., 2015; McNay et al., 2010), especially during aging (Duffy et al., 2019; Freeman et al., 2014).

The genetically tractable fly *Drosophila* is an excellent organism to study the effects of diet on metabolism, behavior, aging and lifespan, due to its short life cycle and the ease by which large number of animals can be reared (Baker and Thummel, 2007; Bellen et al., 2010; Birse et al., 2010; Fontana et al., 2010; Heier and Kühnlein, 2018; Owusu-Ansah and Perrimon, 2014; Padmanabha and Baker, 2014; Rajan and Perrimon, 2013). *Drosophila* has also been employed to model certain human diseases based on the fact that about 65% of human disease-related genes have functional orthologs in flies [see (Ugur et al., 2016; Yamamoto et al., 2014)]. Recently studies have emerged on the effect of HFD on health and physiology of *Drosophila* (Birse et al., 2010; Driver and Cosopodiotis, 1979; Heier and Kühnlein, 2018; Heinrichsen and Haddad, 2012; Huang et al., 2020; Jung et al., 2018; Musselman and Kühnlein, 2018; Rivera et al., 2019; Stobdan et al., 2019; Tatar et al., 2014; Toprak et al., 2020; Trinh and Boulianne, 2013; von Frieling et al., 2020; Woodcock et al., 2015). Most of these studies have investigated effects of relatively short exposure to HFD, ranging between one and three weeks, although some determined the effect of HFD on lifespan (Heinrichsen and Haddad, 2012; Rivera et al., 2019; Woodcock et al., 2015).

It is clear from earlier studies that HFD significantly increases the mortality of both male and female flies and thereby shortens lifespan (Heinrichsen and Haddad, 2012; Rivera et al., 2019; Woodcock et al., 2015), and has detrimental effects on climbing behavior, short-term phototaxis memory, and behavioral responses to odor (Jung et al., 2018; Rivera et al., 2019). Even as short an exposure as one week to HFD has adverse effects on negative geotaxis, heart function, insulin signaling and glucose homeostasis (Birse et al., 2010) as well as gut physiology (von Frieling et al., 2020). Five days of exposure to HFD is sufficient to render flies more sensitive to starvation, seen as an enhancement of the starvation induced hyperactivity (Huang et al., 2020). Furthermore, the HFD was shown to affect the transcription of genes associated with memory, metabolism, olfaction, mitosis, cell signaling, and motor function (Rivera et al., 2019).

The mechanisms by which HFD generates adverse effects on *Drosophila* physiology and longevity are not clear. Lipids are required for energy metabolism, assembly of cellular membranes, biosynthesis of signaling molecules, and in adult female flies especially for oogenesis (Heier and Kühnlein, 2018; Toprak et al., 2020). Since the natural food of *Drosophila* is low in TAG, a main dietary source of stored lipids is carbohydrates, which are converted into TAG in the fat body and intestine (Heier and Kühnlein, 2018). However, many laboratory diets contain lipids [see (Heier and Kühnlein, 2018)], and *Drosophila* has gustatory receptors that sense free fatty acids, FFAs (Ahn and Chen, 2017; Masek and Keene, 2013). These receptors mediate attraction at low levels of FFAs, whereas higher levels lead to repulsion, as monitored in a reflexive feeding response, the proboscis extension reflex (Masek and Keene, 2013). Thus, it appears that flies should be able to regulate their intake of FFAs. Yet, this FFA sensing seems dysregulated or overridden when flies are exposed for extended time to artificial HFD that is based on coconut oil, which consists mainly of FFAs. Thus, flies need to mount a response to minimize the adverse effects of increased intake of dietary FFAs.

We were interested in determining short and long-term effects of HFD on lifespan and physiology in *Drosophila* and found that both virgin and mated female flies exposed to HFD exhibit decreased lifespan, and accelerated decline of climbing ability with age. Virgin females kept for three weeks on HFD display an increased sleep fragmentation compared to controls, and both one and three weeks of HFD results in reduced fecundity. Furthermore, longterm exposure to HFD leads to increased size of the crop, a storage organ associated with the foregut. The distended crops seem to store lipids as determined by Nile Red staining. Surprisingly, we found no long-term effects of HFD on body mass, or levels of triacylglycerids (TAG), glycogen or glucose. However, one week of HFD resulted in decreased body mass and elevated levels of TAG in mated flies.

Mobilization of energy, in the form of stored lipids, during exercise or fasting, is regulated by adipokinetic hormone (AKH), the functional analog of glucagon in insects (Gäde and Auerswald, 2003; Grönke et al., 2007; Heier and Kühnlein, 2018; Toprak et al., 2020; Van der Horst et al., 2001). On the other hand lipogenesis and lipid storage after feeding is regulated by some of the *Drosophila* insulin-like peptides (DILPs) [see (Heier and Kühnlein, 2018; Padmanabha and Baker, 2014; Toprak et al., 2020)]. Thus, we monitored the levels of *Akh* and *dilp* transcripts in flies fed HFD, and found elevated *Akh* transcript, but no effect on *dilp* transcripts. Finally, since *Akh* transcripts are elevated in flies kept on HFD, we investigated the role of AKH in regulating the effects of HFD on lifespan and fecundity. *Akh* mutant flies display increased longevity compared to control flies, but HFD shortens lifespan for both mutants and controls. Also fecundity is affected in *Akh* mutant flies, but short and long term HFD induces similar decreases in fecundity in mutants and controls.

In summary, we could show that HFD activates AKH signaling and has detrimental effects on lifespan, negative geotaxis, sleep and fecundity, as well as induced decreased levels of the biosynthetic enzyme tyrosine hydroxylase in dopaminergic neurons. We also found that the crop may play an important role in the response to HFD.

## 2. Materials and Methods

### 2.1. Fly strains and husbandry

The *Drosophila melanogaster* strain *w^1118^* (from the Bloomington Drosophila Stock Center (BDSC), Bloomington, IN) was used in most experiments. The AKH mutant flies *Akh^AP^* (*w^1118^*; *Akh^AP^*) and their control *w^1118^* (Galikova et al., 2015) were kindly provided by Dr. Ronald Kühnlein (Graz, Austria). This *Akh^AP^* is a null mutation, where also the AKH precursor associated peptide (APRP) is deleted (Galikova et al., 2015). The mutants had been backcrossed to the *w^1118^* line. All the *Drosophila* lines were reared at 25°C, and 12h light/dark cycle. An agar based normal diet (ND) with sugar (10%), yeast (5%) and agar (0.9%) was used for rearing and keeping flies. In all experiments presented in this paper we used female flies. Virgin female flies were obtained by collecting newly hatched flies every three hours and keeping them in vials separate from males.

### 2.2. High-fat diet feeding regime

In all the experiments in this study we used the same HFD food protocol, but two major experimental designs (**Supplemental Fig. 1**). To prepare the high-fat diet (HFD), we added 10% or 30% (volume) of coconut oil to food containing sugar (10%), yeast (5%), and agar (1.2%). Thus, our HFD has the same ratio of sugar (10%), and yeast (5%) as the ND.

To test the effects of HFD on flies, we used two different feeding regimes: (1) the HFD treatment started either within one day of eclosion for virgin females (R1), or (2) 4-5 days after eclosion for mated females (R2). To prevent the flies from getting stuck and drown in the oily HFD food, a piece of moist filter paper was attached to the wall of the food vial, and the food vials were kept in a horizontal position. Flies were sampled at different time points for assays as shown in Supplemental Fig. 1.

### 2.3. Lifespan assay

To test the effects of HFD on lifespan of virgin female flies, one-day-old flies were transferred to vials containing either ND or HFD. Three replicates for each treatment were used for this assay, and for each replicate we used three vials of 15 flies in each. The flies were transferred to fresh food vials every two or three days. Dead flies in each vial were counted and removed when the flies were flipped. The lifespan analysis of *Akh* mutant flies was conducted in the same fashion. Survival curves were generated in Graph Pad Prism version 6.00 (La Jolla, CA) using a log rank Mantel-Cox test.

### 2.4. Negative geotaxis assay and locomotor activity assays

Negative geotaxis (climbing ability) of female flies was measured as described previously (Liao et al., 2017), based on (Gargano et al., 2005). Briefly, flies were kept in upright 35 ml plastic vials. After tapping flies to the bottom of the vial the speed of climbing back was monitored. Flies were allowed 10 seconds to climb back up, at which point a photo was taken. The scoring of climbing ability was performed in two ways, using different flies for the two experiments. (1) The percentage of flies climbing over 5 cm within 10 s was scored (three technical repeats, using six biological replicates) and (2) the height climbed by individual flies was scored from the photo taken after 10 seconds. This was repeated in three technical repeats and the average height each fly had climbed was used for the analysis. Three biological replicates were used for this assay.

The *Drosophila* Activity Monitoring System (DAMS) (Trikinetics, Waltham, MA) was used to analyze locomotor activity and sleep. Virgin female flies were kept on the different diets for three weeks. Thereafter, single flies were placed in glass tubes that in one end had food containing 5% sucrose and 2% agar (sealed with parafilm to prevent desiccation). A small piece of sponge was inserted in the other end of the tube. The tubes with flies were placed into the DAMS within 2 hours before the dark phase. Thus we tracked the activity also during the first night for analysis of individual 24 h periods. For analysis of sleep, locomotor activity was monitored under 12L:12D for 7 days. Only the data collected from the first to the third day were used for the sleep analysis, since flies were transferred to a carbohydrate/ yeast diet and we feared the HFD effect would gradually wear off.

### 2.5. Determination of carbohydrates and triacylglycerids (TAG)

Before measuring the concentration of carbohydrates and TAG, the body weight (wet weight) of single adult flies was determined using a Mettler Toledo MT5 microbalance (Columbus, USA). Whole flies were homogenized in PBS + 0.05% Triton-X 100 using a tissuelyser II from Qiagen Glycogen. Samples were then heat-treated at 70°C for 10 min. After centrifugation at 16000g, 4°C for 10 min, the supernatants were used to determine the concentration of carbohydrates, TAG and protein levels as described below. The supernatants were converted to glucose with 0.5 mg/ml amyloglycosidase from Aspergillus niger (Sigma #10115). Glucose levels were measured with a glucose assay kit with glucose oxidase and peroxidase (Liquick Cor-Glucose diagnostic kit, Cormay, Poland) following the manufacturer’s guidelines. Sample absorbance was measured at 500 nm with a micro-plate reader (Thermo scientific). Glucose concentration was estimated with a linear regression coefficient from a standard curve made from original concentrations of 1.5–15 mg of glucose. Glycogen content was quantified by subtracting free glucose concentration from the total glycogen + glucose levels. The amount of TAG was determined with a Liquick Cor-TG diagnostic kit (Cormay, Poland) using a linear regression coefficient from a standard curve made from an original concentration of 2.2 μg/μl of TAG standard (Cormay, Poland). Sample absorbance was measured at 550 nm with a micro-plate reader (Thermo scientific). Protein levels were determined using a Bradford assay according to Diop and coworkers (Diop et al., 2017). The concentrations of carbohydrates and TAG were normalized to protein levels, and the relative levels were used for generating graphs.

### 2.6. Immunocytochemistry and imaging

*Drosophila* brain, crops, abdominal dorsal vessel (heart) and muscle tissue from flies of different age were dissected out and used for immunostaining. These dissections were performed in the morning for all experiments. The dorsal longitudinal muscles 1d or 1e according to (Fabian et al., 2016) were analyzed in this study. After being dissected out, the tissues were first fixed for 4 h in 4% paraformaldehyde in 0.1 M phosphate buffered saline (PBS) on ice with gently shaking. After washing with PBS, the tissues were incubated with primary antiserum for 48 h at 4°C, followed by four rinses in PBS with 0.5% Triton-X100 (PBS-Tx). Tissues were then incubated with secondary antibody for 48 h at 4°C. After washes in PBS-Tx for 1 hour, the samples were mounted in 80% glycerol in 0.01 M PBS. The following primary antisera were used: Rabbit anti-DILP2 (1:2000) from J.A. Veenstra, Bordeaux, France (Veenstra et al., 2008), rabbit anti-AKH (1:1000) from M.R. Brown, Athens, GA, a monoclonal antibody to mono- and polyubiquitinylated conjugates (FK2, 1:200; from AH diagnostics, Solna, Sweden) [see (Demontis and Perrimon, 2010)], and mouse anti-tyrosine hydroxylase (anti-TH; 1:200) from Incstar Corp., Stillwater, MO, USA (Nässel and Elekes, 1992). Rhodamine-phalloidin (1:1000; Invitrogen) was used to label the muscle fibers of the crop. DAPI (1:2000; Sigma) was used for staining nuclei and Nile red (1:1000; Invitrogen) was used to stain lipids.

As secondary antisera, we used goat anti-rabbit Alexa 546, and goat anti-mouse Alexa 488 antiserum, both from Invitrogen and used at a dilution of 1:1000). All the tissues were scanned with a Zeiss LSM 780 confocal microscope (Jena, Germany) using 10×, 20× or 40× oil immersion objectives. For each experiment, we used identical laser intensities and scan settings. The staining intensity of regions of interest and the nearby background area were measured using Fiji (https://imagej.nih.gov/ij/). Mean staining intensity of the structure was determined by subtracting the background intensity of the same sample. The outline of the crop was extracted manually to determine the crop size (calculated as area).

### 2.7. Capillary feeding (CAFE) assay

The CAFE assay (Ja et al., 2007) was used for measuring food intake and water consumption. Single female flies were placed in 1.5-ml Eppendorf micro centrifuge tubes with an inserted capillary tube (5 μl, Sigma). For the food intake assay, the capillary tube was filled with liquid food containing 5% sucrose, 2% yeast extract and 0.1% propionic acid. This diet, thus, differs from the one used for HFD. To test water consumption relative to feeding, flies were given a capillary tube filled with milli-Q water, in addition to the capillary tube containing food. Eppendorf tubes with food and water capillaries, but without flies, were used as controls for evaporation. The final food intake or water consumption was determined by calculating the decrease in food or water levels minus the average decrease in the control capillaries. Food consumption was measured daily and calculated cumulatively over four consecutive days. We used 8-10 flies in each of three biological replicates.

### 2.8. Fecundity

To determine whether HFD affects fecundity, *w^1118^* mated flies were kept on ND and 30% HFD for either one or three weeks. After the diet treatment, individual females were transferred to ND and the number of eggs laid was counted after 24 h. Three replicates with 19-21 flies in each were tested for each genotype and treatment. To investigate the role of AKH and HFD on fecundity, four-day-old mated *Akh* mutant flies and their *w^1118^* controls were transferred to ND or 30% HFD for one or three weeks. We counted the number of eggs older than stage 10 [according to (Kubrak et al., 2014; Saunders et al., 1989; Shimada et al., 2011) in the ovaries of each fly.

### 2.9. Chill coma recovery

To determine the fitness and stress tolerance of the flies exposed to HFD we tested chill coma recovery. We used *w^1118^* virgin flies exposed to ND, 10% HFD or 30% HFD. A total of 80-94 flies per treatment divided in three biological replicates were used. After one week on the respective diets, groups of four-five flies were placed in empty vials that were placed on ice in a 4°C room. After 130 min on ice, the vials were taken out to room temperature where the behavior of the flies was video recorded for 90 min. The time it took for the flies to recover from the chill coma, defined using righting behavior, was scored from the videos.

### 2.10. Quantitative real-time PCR (qPCR)

Samples for qPCR were taken from virgin female flies. Relative expression of genes of interest and the reference gene ribosome protein 49 (rp49) were determined with qPCR. Total RNA was extracted using Trizol-chloroform (Sigma-Aldrich) from three independent biological replicates. The RNA concentration was measured by NanoDrop 2000 spectrophotometer (Thermo Scientific), the mRNA concentration was adjusted to 400 ng/μl. cDNA was synthesized using random hexamer primer (Thermo Scientific) and RevertAid reverse transcriptase (Thermo Scientific). The cDNA products were diluted 10 times and used for qPCR using a StepOnePlusTM instrument (Applied Biosystem, USA) and SensiFAST SYBR Hi-ROX Kit (Bioline) following the manufacturer’s guidelines. The mRNA levels were normalized to rp49 levels in the same replicate. Relative expression values were determined by the 2^-ΔΔCT^ method (Livak and Schmittgen, 2001). Primer design and evaluation is described in (Kubrak et al., 2014; Kubrak et al., 2016; Liu et al., 2016). The primer sequences used are:

dilp1 F: CGGAAACCACAAACTCTGCG
dilp1 R: CCCAGCAAGCTTTCACGTTT
dilp2 F: AGCAAGCCTTTGTCCTTCATCTC
dilp2 R: ACACCATACTCAGCACCTCGTTG
dilp3 F: TGTGTGTATGGCTTCAACGCAATG
dilp3 R: CACTCAACAGTCTTTCCAGCAGGG;
dilp5 F: GAGGCACCTTGGGCCTATTC
dilp5 R: CATGTGGTGAGATTCGG;
dilp6 F: CCCTTGGCGATGTATTTCCCAACA
dilp6 R: CCGACTTGCAGCACAAATCGGTTA
Akh F: GCGAAGTCCTCATTGCAGCCGT
Akh R: CCAATCCGGCGAGAAGGTCAATTGA
bmm F: GGTCCCTTCAGTCCCTCCTT
bmm R: GCT TGTGAGCATCGTCTGGT
rp49 F: ATCGGTTACGGATCGAACAA
rp49 R: GACAATCTCCTTGCGCTTCT

### 2. 11. Statistics

For statistics and graphs we used Graph Pad Prism version 6.00 (La Jolla, CA, USA2). Results are presented as means ± SEM or means ± range. We initially tested normality of data using Shapiro–Wilk’s normality test. Student’s t-test was used for comparing two groups. For comparing more than two groups, we used one-way analysis of variance (ANOVA) followed by Tukey’s multiple comparisons test, or two-way ANOVA followed by Sidak’s multiple comparisons test. Lifespan data were subjected to survival analysis using a log rank Mantel-Cox test and presented as survival curves. See figure legends for further details.

## 3. Results

We employed two regimes (R1 and R2) for feeding female flies (*w^1118^*) high-fat diet (HFD) (**Supplemental Fig. 1**). The first (R1) was designed according to an earlier study on the effects of diapause on age-related decline in behavior and neuronal senescence (Liao et al., 2017) and used virgin female flies exposed to the experimental diets one day after hatching. The second (R2) was designed to resemble another earlier study (Birse et al., 2010) where mated females were first put on standard diet for several days and then transferred to the experimental diet, but we kept our flies for a longer time on the diet. More specifically in R1 the one-day-old virgin female flies were transferred to vials with standard food (see methods) mixed with 10% or 30% coconut oil (HFD), whereas control flies were kept on normal diet (ND) throughout. The R2 flies were first kept for four days on ND and thereafter the mated female flies were transferred to food enriched with 30% coconut oil (**Supplemental Fig. 1**). This latter protocol allows flies to adjust to adult metabolism on standard food the first four days after hatching. It is known that the first few days after hatching there is residual larval fat body which serves as a nutrient resource while the fly commences feeding and adjusts to its adult metabolism (Aguila et al., 2007). Flies exposed to HFD directly after hatching might thus experience a “dietary stress” during this maturation phase (including ovary maturation). Also, importantly the two regimes allow a comparison between virgin and mated flies in terms of effect of HFD. We first describe data obtained with the virgin flies (R1 protocol).

### 3.1. HFD effects on lifespan, negative geotaxis, metabolism and fecundity

To determine effects of HFD on the physiology, behavior, metabolism and fecundity of female flies we performed a set of assays after different times of exposure to the diet. In some of these we also explored the effect of HFD during ageing of the flies. Since diet composition is critical for egg development and fecundity (Armstrong, 2020; Camus et al., 2019; Hartman et al., 2013; Musselman and Kühnlein, 2018; Partridge et al., 1987), we examined both virgin and mated flies in several of the assays and commence by describing data for virgins.

In tests of lifespan of virgin female flies that were exposed to the HFD food from day one after hatching, both total and median lifespan was shortened with food enriched with 10% lipid, compared to ND controls, and the 30% lipid diet further diminished lifespan (**Fig. 1A**). Median survival times were 33d for controls, 31d for 10% HFD and 28d for 30% HFD, and differences between survival curves for the different diets were significant (P < 0.001).

**Fig. 1.**
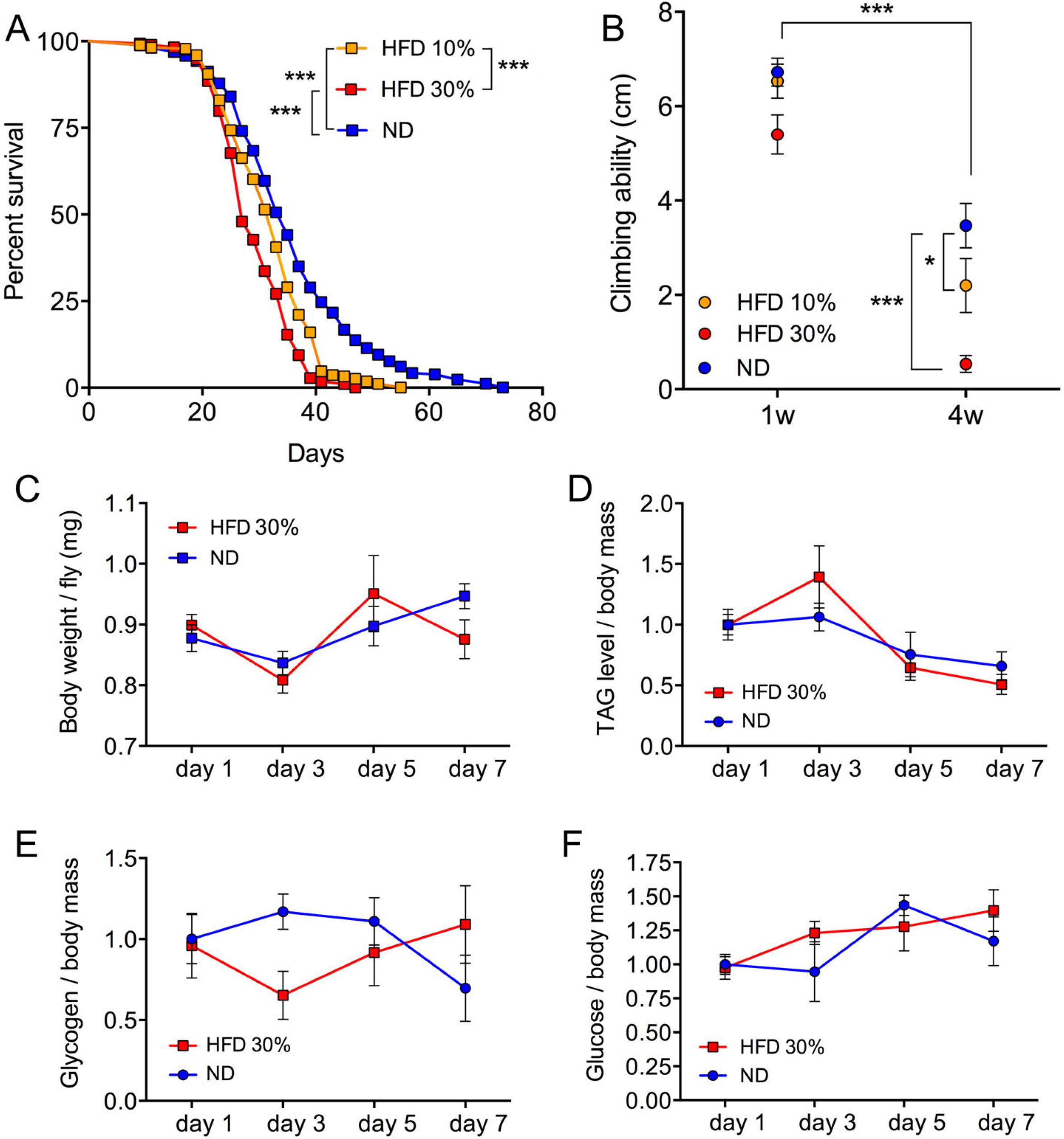
Effects of HFD on lifespan and climbing ability, body weight and levels of TAG, glycogen, and glucose of virgin female flies. **A.** Survival curve of virgin females exposed to ND, 10% HFD and 30% HFD. The survivorship decreased in flies fed with 10% HFD and 30% HFD compared to ND, and the median lifespan is further decreased in flies with 30% HFD compared to 10% HFD. [***p < 0.001 as assessed by log-rank Mantel–Cox test, n = 298–320 flies per treatment, from three independent replicates]. **B.** HFD decreased the climbing ability of four-week-old flies fed 10% and 30% HFD compared to ND. Data are presented as means ± S.E.M, n = 8–19 flies for each group from three independent replicates (*p < 0.05, ***p < 0.001, two-way ANOVA followed by Sidak’s multiple comparisons test). **C - F.** Flies fed with 30% HFD for one to seven days displayed no significant effects on body wet weight, TAG, glycogen and glucose levels compared to ND. Data are presented as means ± S.E.M, n = 6–7 replicates with 5 flies in each replicate for each group (*p < 0.05, **p < 0.01, two-way ANOVA followed Sidak’s multiple comparisons test).

Certain behaviors are known to decline during ageing in *Drosophila* [see (Grotewiel et al., 2005; Tonoki and Davis, 2012; Yamazaki et al., 2014)]. For instance, flies perform progressively worse at negative geotaxis (climbing ability) during normal aging (Gargano et al., 2005). Previous studies showed that exposure to HFD speeds up this deterioration (Jung et al., 2018; Rivera et al., 2019). We performed the climbing assay for virgin flies fed 10% and 30% lipid and noted that after four weeks on the different diets the climbing ability diminished, both for ND and HFD, and that HFD aggravated this deterioration in a dose-dependent manner (**Fig. 1B**). Thus, our HFD protocol yields results similar to published studies.

To determine whether the different diets affected body mass, the flies were weighed at four different time points over one week on the respective diets. Virgin flies exposed to 30% HFD and ND displayed similar body weights at the different time points (**Fig. 1C**). A dose-dependent increase of TAG levels after exposure to HFD has been reported (Birse et al., 2010; Diop et al., 2017; Rivera et al., 2019; Woodcock et al., 2015). We therefore monitored the TAG levels in our virgin flies. To our surprise, the flies on 30% HFD did not differ significantly from those on ND over a week (**Fig. 1D**). Next, we monitored the glucose and glycogen levels, and found that there is no significant difference between HFD and ND (**Fig. 1E and 1F**).

Since the nutrient requirements and metabolism differ between virgin and mated flies, and lipids are important for egg development [see (Armstrong, 2020; Camus et al., 2019; Hartman et al., 2013; Musselman and Kühnlein, 2018; Partridge et al., 1987)], we next went on to investigate mated flies. A comparison of effects of HFD in virgin and mated flies is shown in Table 1. We first determined the body weight and nutrient levels in mated female flies exposed to HFD. In contrast to the virgin flies, mated ones fed 30% HFD displayed decreased body weight compared to ND flies from three days to three weeks of diet treatment (**Fig. 2A**). Also in contrast to the virgin flies, HFD in mated ones has a significant effect on TAG levels with an increase after five days, and then a decrease after three weeks compared to ND (**Fig. 2B**). Glycogen levels do not differ between diets (**Fig. 2C**), whereas glucose levels increase after three days on HFD (**Fig. 2D**). To further compare virgin and mated flies, we also monitored the negative geotaxis of aging mated flies exposed to HFD (**Supplemental Fig. 2A**). At three and four weeks of age, we saw a decline in climbing activity in HFD mated flies, similar to what we found in virgins. However, in older flies (5 w), which were only tested for mated flies, there was no difference between ND and HFD (**Supplemental Fig. 2A**). Thus, to summarize, virgin and mated flies differ somewhat in their metabolic response to HFD (especially TAG levels in mated flies), but display a similar HFD-induced decline in lifespan and performance in the climbing assay with age.

**Fig. 2.**
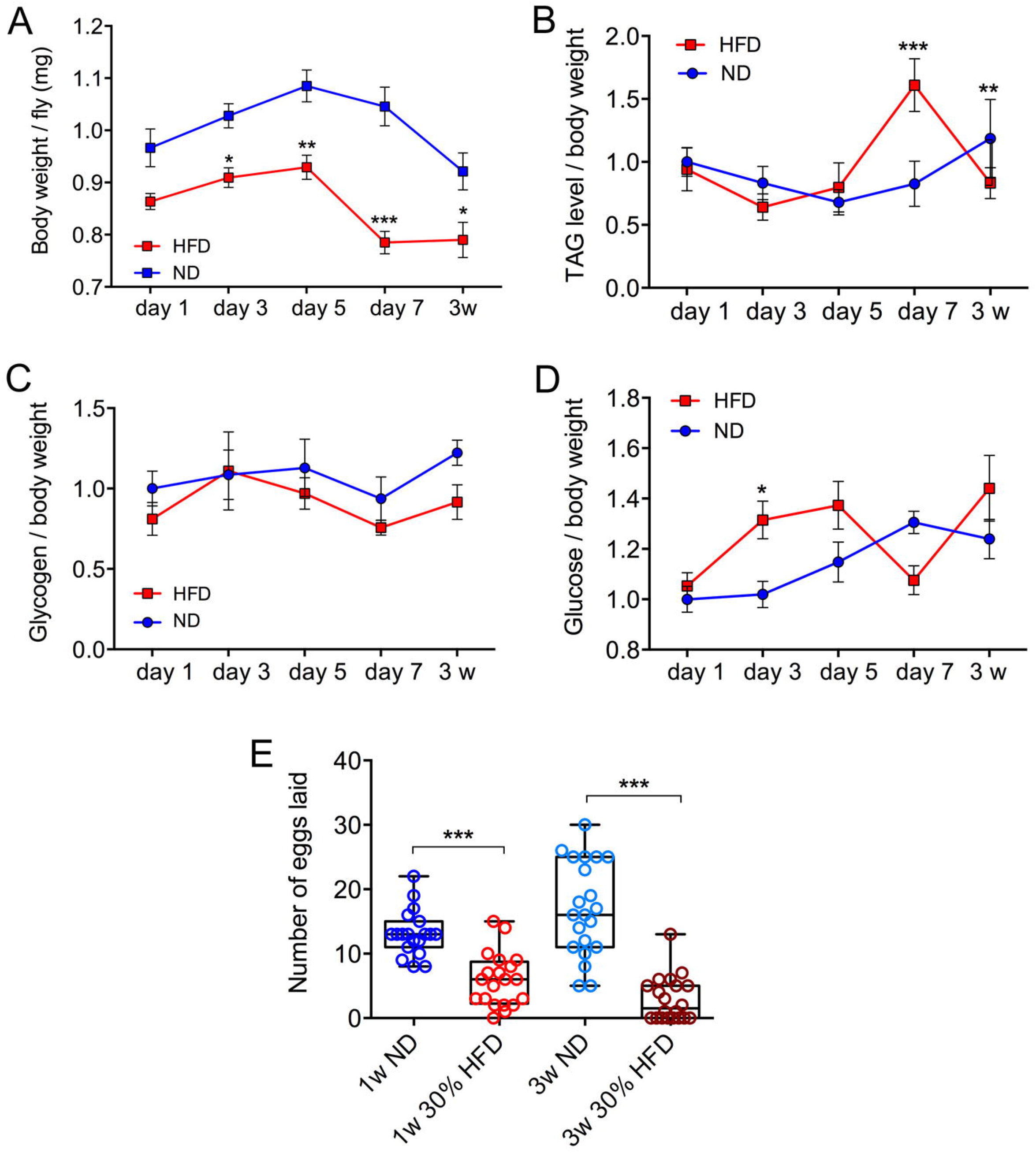
HFD leads to decreased body weight and fecundity, but has no effect on metabolism of mated females. Four-day-old mated flies were fed either ND or 30% HFD. Body wet weight, TAG, glycogen and glucose levels were monitored at day 1, 3, 5, 7, and 3 weeks of age. **A.** Body wet weight is significantly diminished in flies on 30% HFD at all time points after 3 days of age. **B.** The TAG level displays an increase after seven days on HFD, but is lower than in flies on ND after three weeks. **C.** The glycogen level displayed no significant changes with HFD. **D.** The glucose level increased after three days on HFD only. Data in **A-D** are presented as means ± S.E.M, n = 7 replicates with 5 flies in each replicate for each group tested (*p < 0.05, **p < 0.01, **p < 0.001, two-way ANOVA followed by Sidak’s multiple comparisons test). **F.** Mated flies kept on 30% HFD for one or three weeks laid significantly fewer eggs compared to flies kept on ND. Data are presented as means ± S.E.M, n = 19–21 flies for each group from three independent replicates (***p < 0.001, one-way ANOVA followed by Tukey’s multiple comparisons).

**Table 1.** Effects of high fat diet (30%) on virgin and mated flies each compared to normal diet

| Assay | Virgin female flies |  |  | Mated female flies |  |  |
| --- | --- | --- | --- | --- | --- | --- |
|  | Effect | Significance | Fig. | Effect | Significance | Fig. |
| Lifespan | decrease | *** | 1A | decrease | *** | 6J |
| Climbing<sup>1</sup> | decline | *** | 1B | decline | * | S 2A |
| Weight<sup>2</sup> | no change | ns | 1C | decrease | *** | 2A |
| TAG<sup>2</sup> | no change | ns | 1D | increase | *** | 2B |
| Glycogen<sup>2</sup> | no change | ns | 1E | no change | ns | 2C |
| Glucose<sup>2</sup> | no change | ns | 1F | increase<sup>4</sup> | * | 2D |
| Fecundity<sup>3</sup> | nt |  |  | decrease | *** | 2E, 7 |
| Sleep bouts<sup>5</sup> | increase | ** | 3F | nt |  |  |
| Crop size<sup>5</sup> | nt |  |  | increase | *** | 4A-H, 8 |
| Food intake<sup>6</sup> | nt |  |  | increase | ** *** | 5A,B |
| Water intake<sup>6</sup> | nt |  |  | decrease | ** | 5C |
| *Akh* transcript | increase | * | 6A | nt |  |  |

**Notes:**
nt, not tested
<sup>1</sup> Four weeks on HFD
<sup>2</sup> Seven days on HFD for virgins. Mated flies HFD for seven days and three weeks. TAG increase seen in mated flies after seven days
<sup>3</sup> One and three weeks on HFD, monitored as eggs remaining in fly and eggs laid
<sup>4</sup> Increase was only seen after three days on HFD
<sup>5</sup> After three weeks on HFD
<sup>6</sup> In 5A determined by CAFE assay after 7 days on HFD; in 5B after 7 days on HFD, flies were given a choice between water and food for 24h
Significance: \*p < 0.05, \*\*p < 0.01, \*\*\*p < 0.001 (see figure legends for further data)

Cold tolerance in *Drosophila* depends on many factors including regulation of ion homeostasis, as well as carbohydrate and lipid metabolism [see (Overgaard and MacMillan, 2017; Sinclair and Marshall, 2018)]. Insects, including *Drosophila*, respond to prolonged cold by entering a coma-like state and after transfer to normal temperature they recover (Overgaard and MacMillan, 2017; Sinclair et al., 2013). The time it takes to recover from chill coma is used as a measure of cold tolerance (MacMillan et al., 2018; Overgaard and MacMillan, 2017; Terhzaz et al., 2015). To test whether HFD affects cold tolerance, we monitored the recovery time from chill coma in virgin female flies after eight days of exposure to 10% or 30% HFD and ND (**Supplemental Fig. 2B**). The 30% HFD flies displayed significantly faster recovery from chill coma, suggesting that short-term exposure to HFD is beneficial for cold tolerance, at least in virgin flies.

Fecundity, including egg development, in *Drosophila* females depends on nutrient availability and diet composition (Armstrong, 2020; Camus et al., 2019; Hartman et al., 2013; Musselman and Kühnlein, 2018; Partridge et al., 1987). Therefore, we asked whether the fecundity of the mated flies is affected by HFD. We exposed one- and three-week-old mated flies to 30% HFD or ND for one or three weeks and then monitored the number of eggs laid over 24 h on plates with ND food (since HFD is too sticky). Both after one and three weeks, the HFD results in a significantly decreased number of eggs laid (**Fig. 2E**). It cannot be excluded that the transfer of HFD flies to ND for 24h of egg laying affects egg numbers somewhat.

### 3.2. HFD during ageing affects activity and sleep

Both locomotor activity and sleep patterns change with ageing in *Drosophila* (Liao et al., 2017; Luo et al., 2012). Since we could show that HFD affects climbing performance during ageing, we also tested the long-term effects of HFD on locomotor activity and sleep. For this, we only monitored virgin flies since they display a stronger biphasic daily activity than mated ones [see (Isaac et al., 2010)]. The flies (R1 protocol) were kept on the different diets (ND, 10% HFD or 30% HFD) for one or three weeks, where after locomotor activity and sleep were monitored using the *Drosophila* Activity Monitor System (DAMS, Trikinetics). Note that after transfer to the DAMS all the flies received 5% sucrose (in agar) in the capillary tubes since the HFD is too oily and flies would get stuck. The average activity histograms across the first three days are shown in **Fig. 3A and 3B**. Note that during the first night after transfer to the DAMS, the flies kept for three weeks in 30% HFD exhibited hyperactivity (**Fig. 3B**, arrow). This is consistent with a recent study that demonstrated that HFD enhanced starvation induced hyperactivity (Huang et al., 2020). Next, we analyzed the average activity (using 5 min bins) of flies fed ND, 10% HFD and 30% HFD for one or three weeks. The 30% HFD flies (both after one and three weeks of diet) displayed less activity than ND flies during the light period, but 10% HFD did not affect activity (**Supplemental Fig. 3A-D**). The average total activity during 24 h calculated over three days indicates that the 30% HFD flies are less active than the ND flies both after one and three weeks of feeding the diet (**Supplemental Fig. 3 E**).

**Fig. 3.**
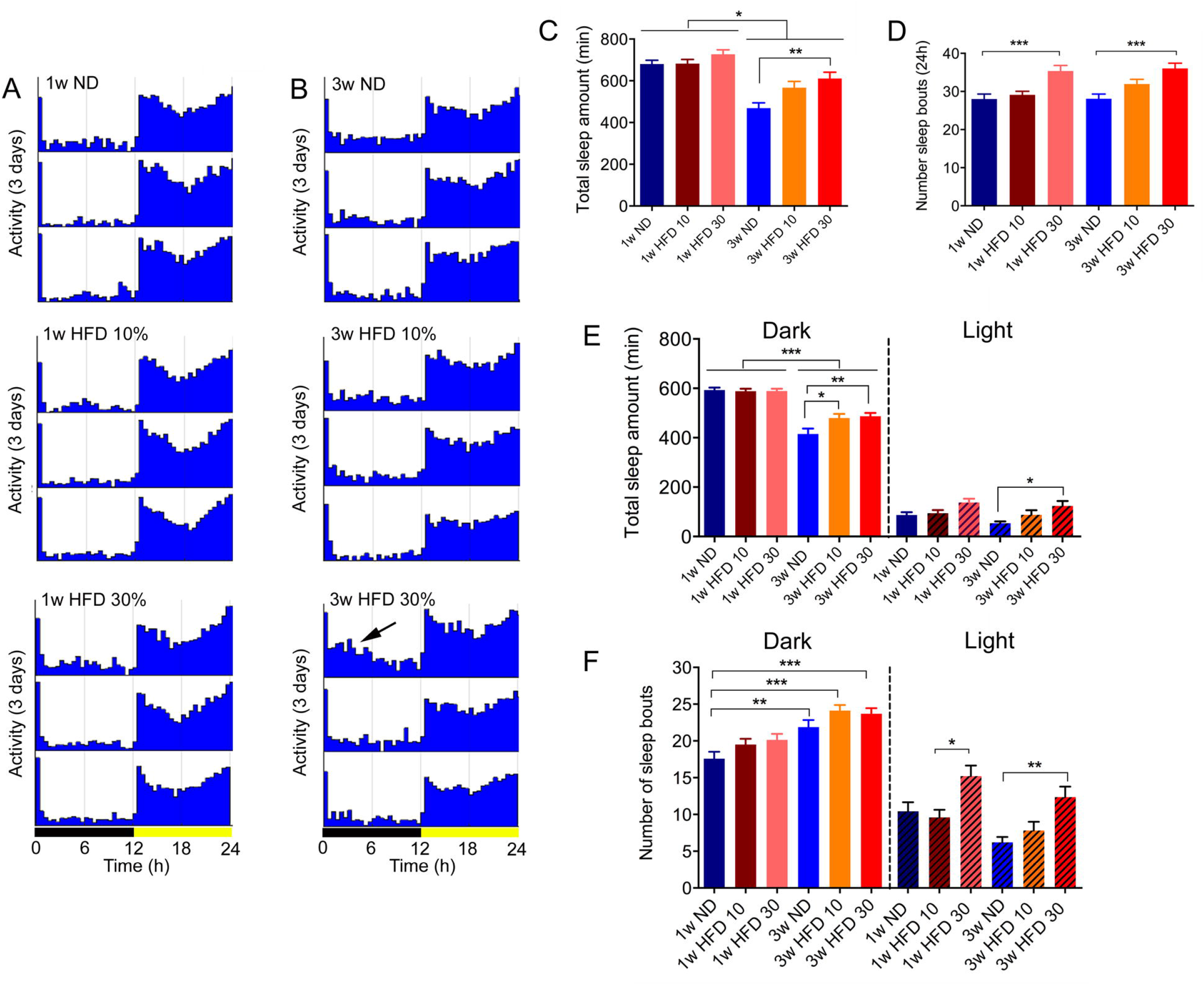
HFD leads to hyperactivity and sleep fragmentation in virgin female flies. **A.** Average activity histograms over three days during light and dark phase from flies fed with ND, 10% HFD and 30% HFD for one week. Activity monitoring after HFD was performed on flies that obtained food containing 5% sucrose in their capillaries. **B.** Activity histograms from flies fed with ND, 10% HFD and 30% HFD for three weeks (otherwise as in A). Note that the flies fed HFD for 3 weeks before activity monitoring displayed increased activity during the first night (arrow). **C.** The total sleep time is decreased in three-week-old flies compared to one-week-old flies, and flies fed 30% HFD for 3 weeks displayed increased amount of sleep compared to ND. **D.** Average number of sleep bouts per 24 h. The number of sleep bouts averaged over three 24 hour periods is higher in flies exposed to 30% HFD both after 1 week and 3 weeks. **E.** The total sleep time shown separately for the light and dark phases. Aging decreased sleep time during dark phase. Total sleep time decreased during the dark phase in aging flies, and flies fed HFD for 3 weeks showed increased sleep time during both the light and dark phases. **F.** The number of sleep bouts during light and dark phases of flies fed with ND and HFD for 1 and 3 weeks. HFD and aging increase the number of sleep bouts. During the light period 30% HFD increased sleep bouts both after one and three weeks of the diet. Data in graphs are presented as means ± S.E.M, n = 46-48 flies for each treatment from three replicates (*p < 0.05, **p < 0.01, ***p < 0.001, as assessed by one-way ANOVA followed by Tukey’s multiple comparisons).

Since ageing impairs sleep patterns in *Drosophila* (Luo et al., 2012; Umezaki et al., 2012), we next monitored the total amount of sleep and the number of sleep bouts over 24 h in virgin flies kept for one or three weeks on the diets. Regardless of diet, the total amount of sleep decreased with age (**Fig. 3C**). After three weeks of 30% HFD flies slept more than the ND flies, whereas flies on 10% HFD are not significantly different from ND or 30% HFD (**Fig. 3C**). The number of sleep bouts (day and night; 24 h) is significantly higher in 30% HFD flies compared to ND fed flies both after one week and after three weeks (**Fig. 3D**). When we break up the sleep data into daytime and nighttime sleep (light and dark period), we see that the total nighttime sleep decreases with age and that the flies fed 10% and 30% HFD sleep more than ND flies (**Fig. 3E**). The daytime sleep does not change with age, but flies kept three weeks on the 30% HFD diet sleep more than the ND flies (**Fig. 3E**). During nighttime, the number of sleep bouts increased after three weeks, compared to one week in all three diets (**Fig. 3F**). This indicates an increase in sleep fragmentation during aging, as described before (Liao et al., 2017; Metaxakis et al., 2014; Williams et al., 2016). During daytime, the 30% HFD flies display a higher number of sleep bouts compared to both ND at 3 weeks and compared to 10% HFD flies at one week (**Fig. 3F**). This suggests that 30% HFD impairs sleep quality by increasing sleep fragmentation, but the effect of ageing does not worsen after one week.

### 3.3. HFD effects on aging of neurons and muscle

Next, we asked whether it is possible to identify cellular changes that correlate with the behavioral senescence caused by HFD in aging flies, and chose to analyze dopaminergic neurons, skeletal and heart muscle. The dopaminergic system in *Drosophila* plays a critical role in learning and memory, as well as in regulating locomotor behavior, aggression, arousal, food search, and stress (Alekseyenko et al., 2013; Aso and Rubin, 2016; Ichinose et al., 2017; Landayan et al., 2018; Petruccelli et al., 2020; Riemensperger et al., 2013; White et al., 2010). It has been shown that dopamine (DA) levels decrease with age in brain neurons of *Drosophila* (Liao et al., 2017; Neckameyer et al., 2000; White et al., 2010). This was demonstrated by immunocytochemistry with antiserum to the rate limiting biosynthetic enzyme tyrosine hydroxylase (TH). Here, we monitored the TH-immunoreactivity in a cluster of brain DA neurons of flies kept for one or three weeks on ND and three weeks of 30% HFD, using virgin flies (**Supplemental Fig. 4A, B**). Analyzing the protocerebral posterior lateral (PPL1) cluster of DA neurons (Nässel and Elekes, 1992) we found that there is no significant difference in TH immunolabeling in PPL1 neurons after three weeks of aging under ND conditions (**Supplemental Fig. 4A, B**). However, three weeks of 30% HFD led to strongly decreased TH levels in PPL1 neurons compared to one and three weeks of ND (**Supplemental Fig. 4A, B**). It can be noted that earlier studies found decreased TH levels after eight weeks of ND [see (Liao et al., 2017)].

Since deterioration of muscle function with ageing might affect climbing and locomotor activity (Beramendi et al., 2007; Liao et al., 2017), we next monitored muscle of flies fed different diets. One marker for ageing in muscle fibers of flies is an increased deposition of aggregates containing polyubiquitinated proteins (Demontis and Perrimon, 2010). We compared the anti-polyubiquitin immunoreactivity levels in abdominal muscles of normally aging virgin flies and flies fed 30% HFD (**Supplemental Fig. 4C-F**). Our data show an increase in number and size of polyubiquitin particles over five weeks for both ND and HFD, but HFD in itself has no significant effect on the accumulation of polyubiquitinated particles in these muscle compared to flies kept on ND (**Supplemental Fig. 4C-F**). It is known that short-term exposure to HFD has an adverse affect on the morphology of the heart, also known as abdominal dorsal vessel (Birse et al., 2010), we also analyzed the heart using phalloidin staining to visualize muscle fibers. Surprisingly, we detected no effect of exposing virgin flies to HFD for one or five weeks compared to flies kept on ND (**Supplemental Fig. 4G**). However, the hearts of five-week-old flies on both diets displayed more compact muscle fibers.

### 3.4. HFD effects on crop size and food ingestion

Adult flies use the crop, a diverticulum from the foregut, for temporally storing nutrients, primarily carbohydrates (Stoffolano and Haselton, 2013). In our experiments, we observed that the crop is dramatically enlarged in mated flies kept on HFD for three weeks compared to ND flies (**Fig. 4A-H**). Using Nile red to stain lipids, we found that the enlarged crop is filled with aggregates of lipid (**Fig. 4F, G**), which was not seen in flies kept on ND (**Fig. 4D, E**). These findings suggest that the HFD affects the emptying of the crop and that it stores lipids, presumably free fatty acids (FFAs) of the coconut oil.

**Fig. 4.**
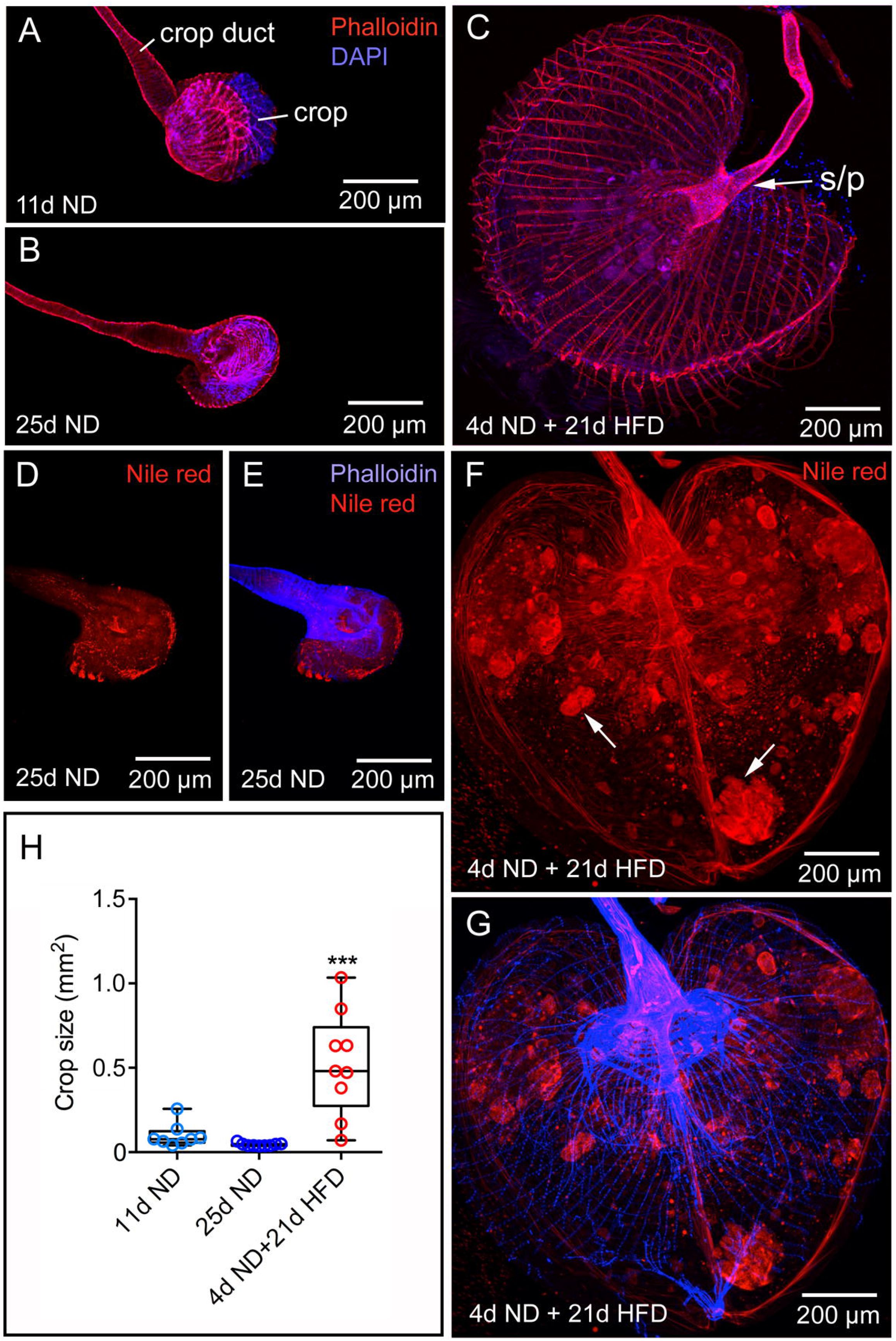
The crop is enlarged in mated female flies fed HFD. Flies were kept on ND for 11 days, 25 days or transferred to 30% HFD at 4 days of age whereafter they were kept on the HFD for 21 days (reaching a total age of 25 days). The morphology of the crops was investigated after staining with rhodamine-phalloidin (red) and DAPI (blue). **A.** The morphology of the crop and crop duct of flies kept for 11 days on ND. **B.** Crop from a fly kept for 25 days on ND. **C.** Flies kept for 4 days on ND and then 21 days on 30% HFD displayed a drastically enlarged crop. s/p, sphincter and pump region [one of several, see (Stoffolano and Haselton, 2013)]. **D** and **E**. Nile red staining of crop from control (fly fed for 25 d on normal diet). Muscle fibers are stained with rhodamine-phalloidin (blue). **F** and **G.** Nile Red staining reveals that the crop of a fly fed with HFD for 21 days is loaded with lipid aggregates (e. g. at arrows). Rhodamine-phalloidin shown in blue. **H.** Size quantification (total area) of fly crops after 11 days ND, 25 days ND and 4 days ND plus 21 days HFD. Data are presented as means ± S.E.M, n = 8-9 flies for each treatment from three replicates (***p < 0.001, compared to one-week-old ND flies, as assessed by one way ANOVA followed by Tukey’s multiple comparisons).

Recently it was shown that the crop is providing feedback to the brain to regulate food ingestion (Wang et al., 2020). These authors found that proprioceptors receptors (Piezo) in the crop that monitor crop filling signal to neurons in the brain that regulate feeding. Since the crop size increased in flies fed HFD, we asked whether HFD would affect food ingestion. Food consumption (using ND food) was measured in mated flies using the CAFE assay. Note that the ND food given to both experimental groups in the CAFE assay is based on 5% sucrose and 2% yeast extract; thus, HFD flies experience a switch in diet. This was intentional since we wanted to determine whether the HFD flies consumed more or less ND food, without bias for sensing of dietary free fatty acids in the coconut oil by specific gustatory neurons (Ahn and Chen, 2017; Masek and Keene, 2013). We found that flies fed HFD for seven days ingest significantly more standard food (ND) during the first day after transfer to the CAFE assay than control flies kept for the same time on ND (**Fig. 5A**). Thus, it is possible that the crop distension is monitored by Piezo-expressing neurons and relayed to brain circuits that diminish food intake in the short term.

**Fig. 5.**
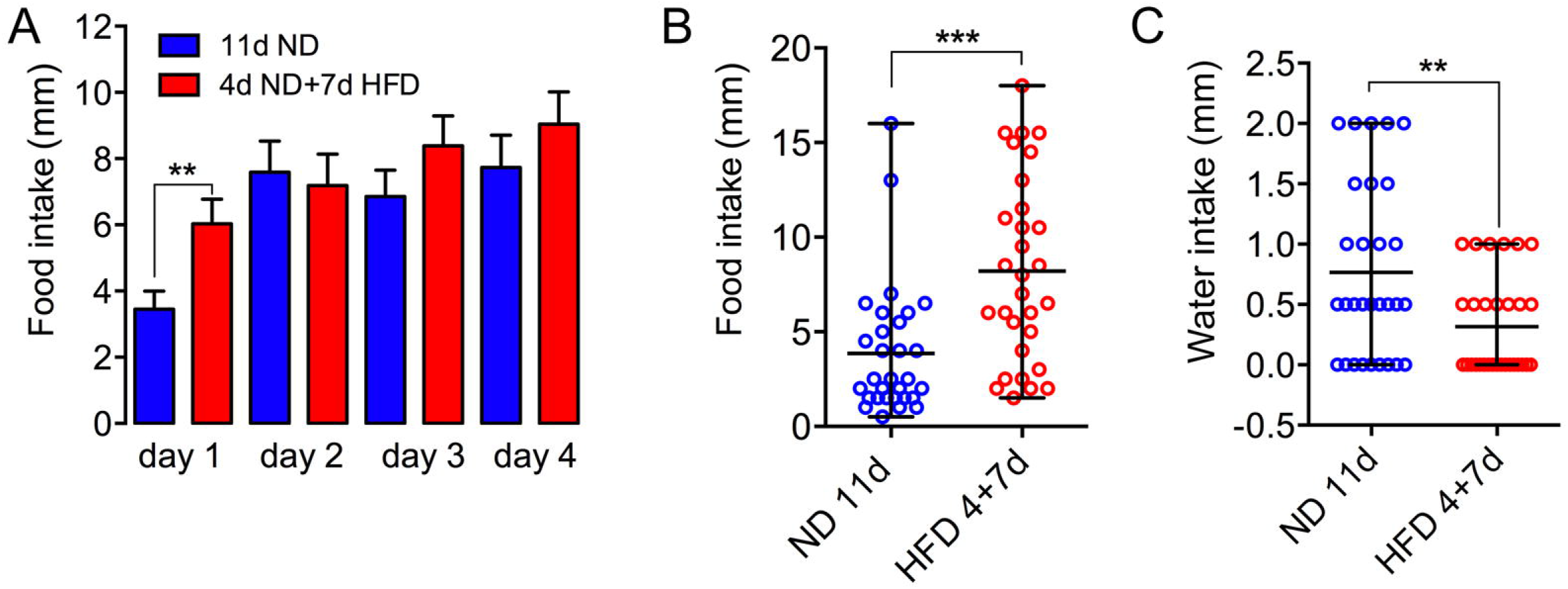
HFD affects feeding and drinking. **A.** Food intake increased in mated flies kept for seven days on HFD only during the first day after transfer to the CAFE assay (where ND was provided). Data are presented as mean ± S.E.M, n = 24-30 flies for each treatment from three replicates. (**p < 0.01, unpaired Student’s t-test). **B and C.** When given a choice between food and water for 24h in the CAFE assay, mated flies kept for 7 days on 30% HFD consumed more food (**B**) but less water (**C**) compared to ND flies. Data are presented as mean ± range, n = 30 flies for each treatment from three replicates (**p < 0.01, ***p < 0.001, unpaired Student’s t-test).

Since flies kept in the CAFE assay obtain no separate water for drinking (they only obtain water available in the food), we next tested whether the increased food intake is due to thirst or hunger. Thus, we exposed the mated flies to a food and water choice experiment. In a CAFE assay set-up, we inserted one capillary tube with water and another tube containing food (ND). Flies kept for seven days on 30% HFD were monitored for 24 h. Our data show that the flies kept on HFD consume more food and drink less water than flies kept on ND (**Fig. 5B, C**). These findings suggest that flies kept on HFD experience hunger, at least in the short-term.

### 3.5. Effects of HFD on hormonal signaling

Lipid and carbohydrate storage and metabolism are regulated hormonally. Among the key hormones are adipokinetic hormone (AKH), the functional analog of mammalian glucagon, (Bharucha et al., 2008; Grönke et al., 2007; Kim and Rulifson, 2004; Lee and Park, 2004) and *Drosophila* insulin-like peptides (DILPs) (Broughton et al., 2005; Grönke et al., 2010; Rulifson et al., 2002; Zhang et al., 2009). Thus, we investigated the effects of HFD on transcript levels of *Akh, dilp1*, *dilp2, dilp3, dilp5,* and *dilp6.* Virgin flies were kept on ND or 30% HFD for one or five weeks to test effects of diet and ageing. At these time points, the transcript levels of several genes of interest were determined by qPCR. The *Akh* transcript level is four-fold higher in the five-week-old HFD flies compared to those kept for one week on either diet, and compared to five-week-old flies on ND (**Fig. 6A**). The lipase Brummer (encoded on *bmm*), which is a homolog of the mammalian adipose triglyceride lipase, is critical for lipid metabolism and mobilization in *Drosophila* (Grönke et al., 2007). We found that *bmm* levels increased significantly only in flies kept five weeks on ND with no additional effect of HFD (**Fig. 6B**). Levels of *dilp1, dilp3, dilp5,* and *dilp6* were not found significantly different between diets and no effects of ageing were observed, with the exception of a slight increase in *dilp2* transcript after 5 weeks on ND (**Fig. 6C-G**). Moreover, we monitored the AKH peptide levels by immunohistochemistry in the adult corpora cardiaca, and found no changes in AKH levels with diet or aging (**Fig. 6H**). Taken together with the elevated *Akh* transcript, this might suggest that AKH production and release increases as a consequence of HFD.

**Fig. 6.**
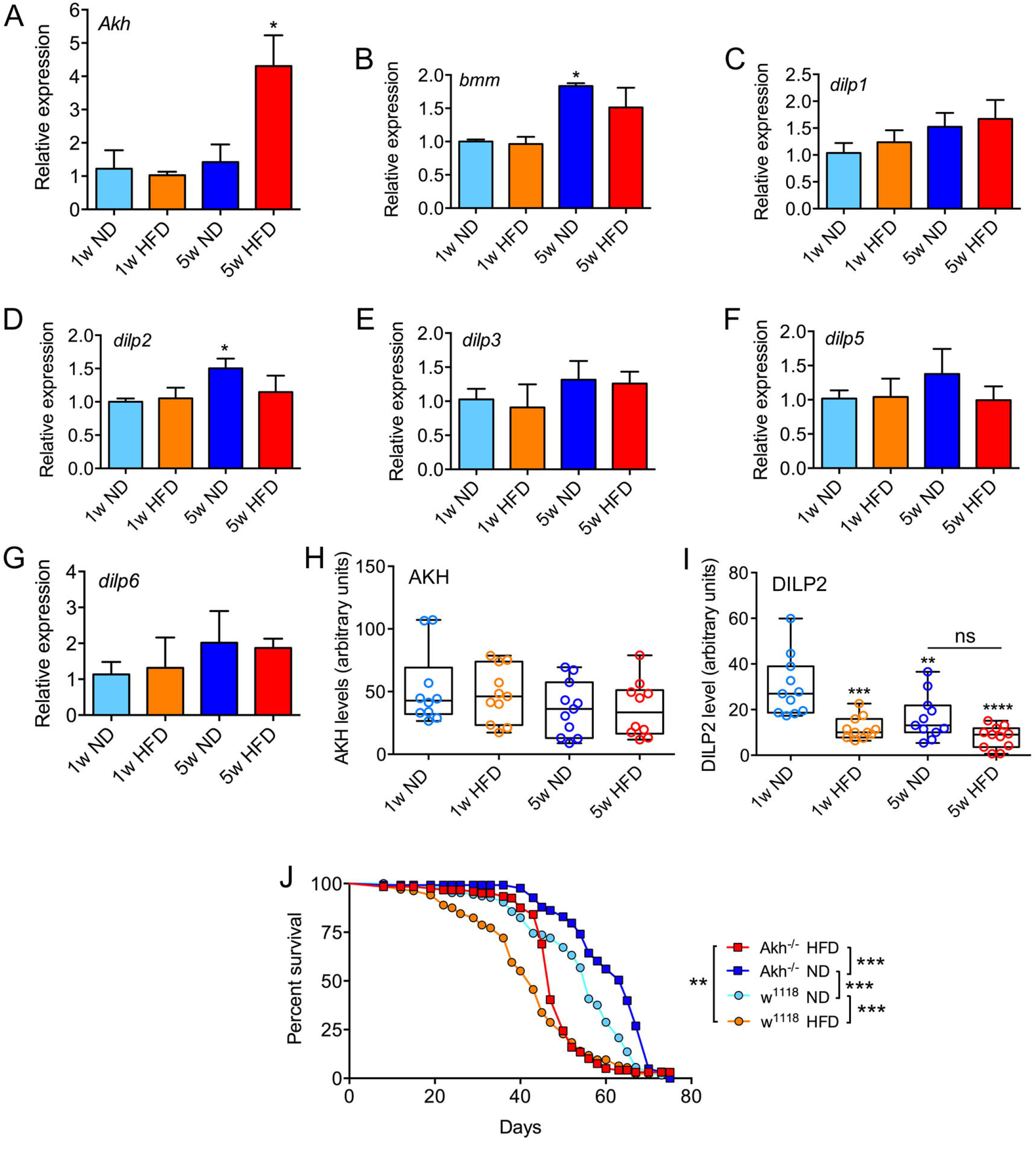
The effects of aging and HFD influence transcripts of genes related to lipid metabolism and lifespan of mutant flies. One-day-old virgin female flies were transferred to ND or 30% HFD. Gene expression and protein levels were monitored after 1 and 5 weeks on the diets. **A.** *Akh* expression increased after 5 weeks on HFD. **B.** A slight increase in *bmm* levels is seen after 5 weeks on ND, but no significant effect of HFD. **C-G.** The levels of different *dilp* transcripts do not change with HFD. A slight increase is seen for *dilp2* in five-week-old flies on ND. The data in A-G are presented as means ± S.E.M.; three replicates with 10 flies in each replicates were analyzed (*p < 0.05, as assessed by one-way ANOVA followed by Tukey’s multiple comparisons). **H.** AKH immunolabeling in corpora cardiaca does not change with age or diet. Data are presented as means ± S.E.M, n = 10-11 flies from three replicates for each treatment. Differences among groups were assessed by one-way ANOVA followed by Tukey’s multiple comparisons). **I.** DILP2 immunolabeling decreased after one week on HFD compared to ND. After 5 weeks, a decrease in DILP2 levels was observed in both treatment groups compared to one-week-old flies kept on ND. However, after 5 weeks no difference is seen between ND and HFD. Note that the asterisks indicate significance levels compared to one week on ND. Data are presented as means ± S.E.M, n = 11 flies for each treatment from three replicates (**p < 0.01, ***p < 0.001, as assessed by one way ANOVA followed by Tukey’s multiple comparisons). **J.** Survival curves of virgin *Akh* mutants and their control *w^1118^* under ND and HFD. *Akh* mutant flies display increased median lifespan compared to *w^1118^* flies under both ND and HFD conditions and HFD diminished the median life span of both *Akh* mutant and *w^1118^* flies. Median lifespans: for the genotypes are *w^1118^* on ND: 56d, *w^1118^* on HFD: 43d, *Akh^-/-^* on ND: 65d, *Akh^-/-^* on HFD: 47d (***p < 0.001 for all comparisons as assessed by log-rank Mantel–Cox test, n = 124–136 flies per treatment from three independent replicates).

In contrast, the DILP2 peptide levels in the insulin producing cells (IPCs) were reduced as an effect of both diet and age. Flies fed HFD for one week exhibited much lower DILP2 levels compared to ND flies (**Fig. 6I**). This suggests that short-term exposure to HFD may increase DILP2 release from IPCs. Five weeks of exposure to both ND and HFD led to decreased DILP2 levels compared to younger flies, suggesting an impact of ageing, but no effect of long term exposure to HFD (**Fig. 6I**). A diminished DILP2 level with ageing was also noted in a previous study (Liao et al., 2017).

### 3.6. AKH signaling and effects of HFD on lifespan, fecundity and crop size

Increased AKH signaling, resulting from various genetic interventions, has been shown to extend lifespan in *Drosophila* (Post et al., 2019; Waterson et al., 2014). Thus, we next asked whether lifespan is affected by HFD when AKH signaling is disrupted. To do this, we examined the survival of *Akh* null mutant flies *[w^1118^;Akh^AP^;* (Galikova et al., 2015)], kept on ND and HFD. Interestingly, we found that under both ND and HFD conditions, virgin *Akh* mutants display significantly increased median lifespan compared to control flies (*w^1118^*) on the same diets (**Fig. 6J**). Hence, in our experiments, diminished AKH signaling increases survival of the flies, regardless of diet. Furthermore, HFD shortens the median lifespan of both *Akh* mutants and *w^1118^* flies compared to the same genotypes on ND (**Fig. 6J**).

It has been proposed that AKH is important in mobilizing energy to support oocyte growth in insects (Lorenz and Gäde, 2009). We therefore asked whether HFD and AKH signaling have an effect on oogenesis. Thus, we exposed four-day-old mated *Akh* mutant and *w^1118^* flies to ND and HFD for one week and monitored the number of mature eggs in their ovaries. We found that under ND conditions, the ovaries of *Akh* mutant flies contained much fewer eggs than controls (**Fig. 7A**). This supports the importance of AKH signaling for lipid mobilization and oogenesis. Moreover, the number of eggs is drastically lower in flies exposed to one week of HFD for both the *Akh* mutants and *w^1118^* (**Fig. 7A**), suggesting that HFD is detrimental for fecundity. Next, we also analyzed *Akh* mutant and *w^1118^* flies exposed to ND and HFD for three weeks (**Fig. 7B**). The results are similar to those from one-week exposure, except that on ND ovaries of *Akh* mutant flies contains a larger number of eggs than controls. However, HFD leads to decreased number of eggs in both mutants and controls. These findings of reduced egg numbers in ovaries of HFD *w^1118^* flies (**Fig. 7A, B**), combined with our data in section 3.1 showing that HFD results in reduced egg laying (**Fig. 2G**), suggest that HFD diminishes fecundity. In the short term deletion of AKH signaling impairs fecundity in flies on ND, whereas in the longer term it seems beneficial. An earlier study using the same *Akh^AP^* mutant flies found no effect on fecundity as calculated as daily scores of eggs laid over ten days (Galikova et al., 2015).

**Fig. 7.**
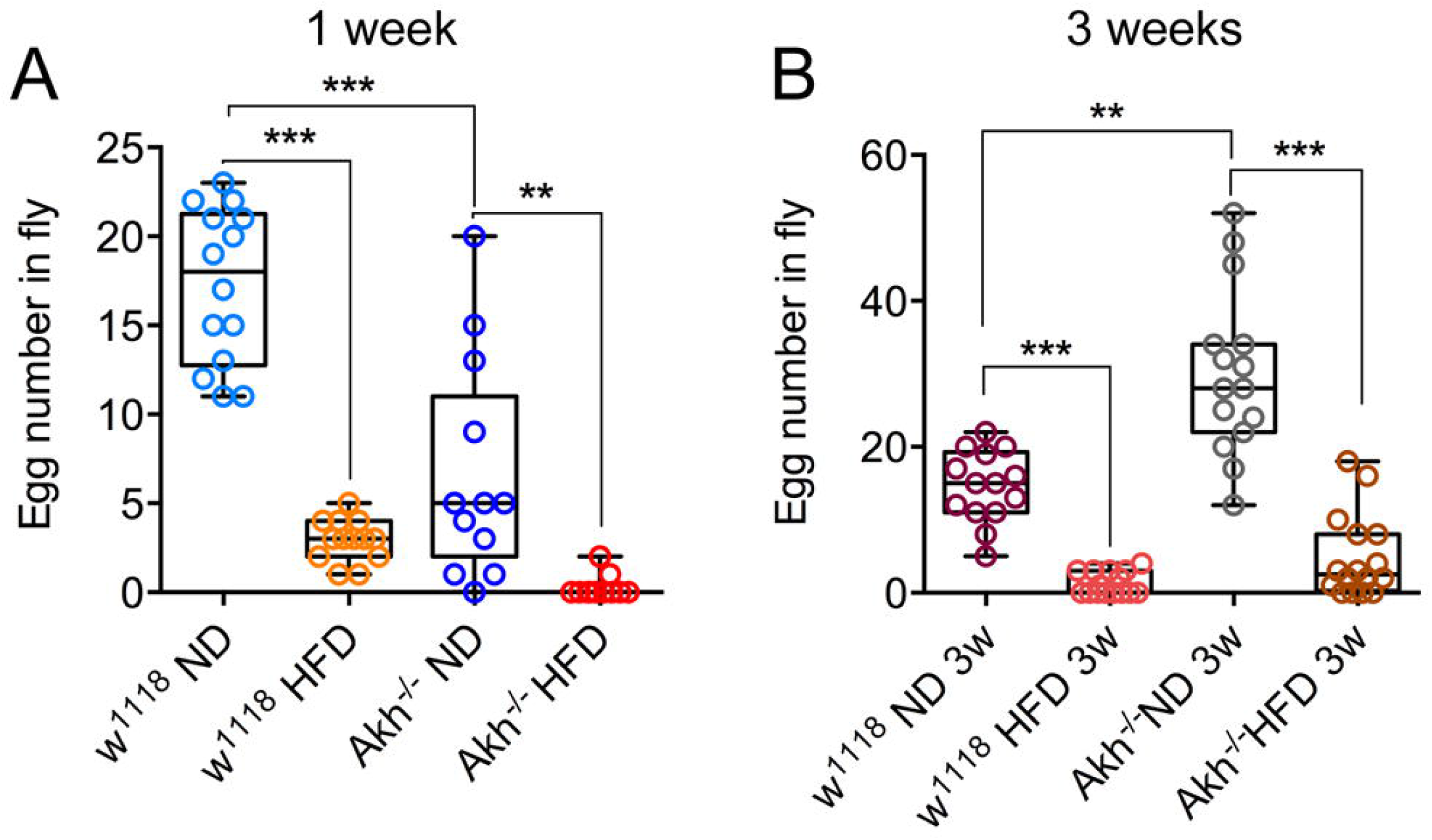
HFD for one or three weeks affects fecundity. **A.** HFD for one week decreased egg numbers in ovaries of mated flies compared to ND for both *Akh* mutants and *w^1118^* controls, and *Akh* mutants have fewer eggs compared to *w^1118^* controls. **B.** Also after three weeks on HFD egg numbers are decreased in both *Akh* mutants and controls compared to ND. However, in *Akh* mutant flies three weeks on ND results in significantly more eggs than in control flies on the same diet. Data are presented as means ± S.E.M, n = 11-14 flies for each treatment from three replicates (**p < 0.01, ***p < 0.001, as assessed by one way ANOVA followed by Tukey’s multiple comparisons).

We showed in section 3.4 above that exposure to HFD for three weeks leads to an enlargement of the crop (**Fig. 4**). The size of the crop is under neuronal and hormonal regulation (Gough et al., 2017; Kuraishi et al., 2015; Solari et al., 2017). Interestingly, AKH stimulates increases in the frequency of contractions of muscle fibers of the crop, including those in crop pumps that are involved in moving food contents (Solari et al., 2017). Thus, we monitored the crop size in mated *Akh* mutants and *w^1118^* controls exposed to the two diets. As seen in **Figure 8**, three weeks of HFD induce the same increase in crop size in mutants and controls, suggesting that in the long term AKH signaling makes little difference for crop volume.

**Fig. 8.**
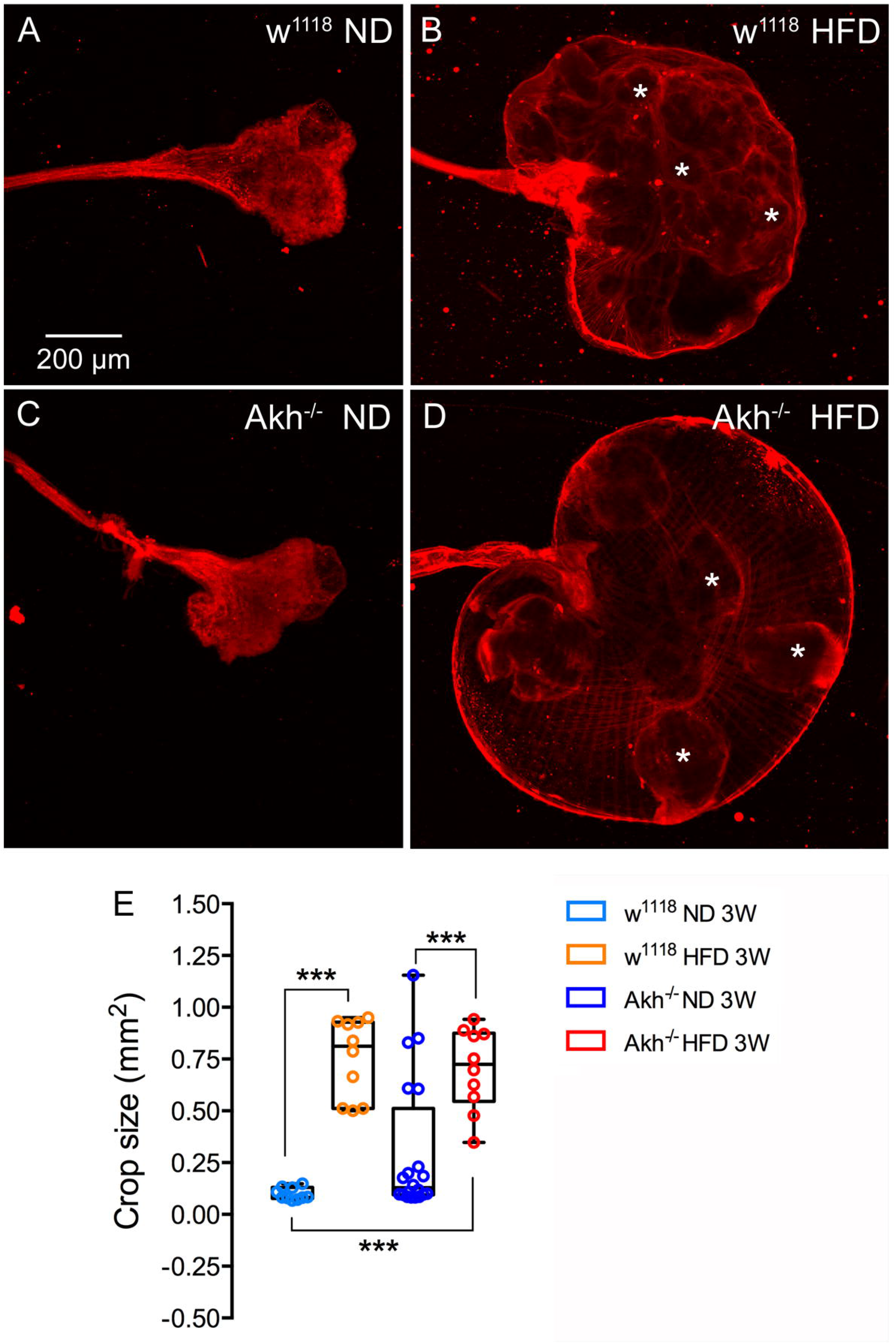
The effect of HFD on the size of the crop is similar in mated *Akh* mutant flies and controls. **A** and **B**. In control flies (w^1118^) the crop size increases after three weeks on high fat diet (HFD) compared to normal diet (ND). Nile red staining shows that the HFD results in large lipid aggregates in the crop (e. g. at asterisks). **C** and **D**. In AKH mutant flies (Akh^-/-^) crop size also increases with HFD. **E**. Quantification of crop size. HFD results in significantly increased crop size both in controls and mutants. There is no significant difference between *Akh* mutants and controls kept on ND or on HFD. However, note the larger variability in crop size of mutants kept on normal diet. These enlarged ND crops do not contain Nile red stained lipid aggregates (see Supplementary Figure 5C). Data are presented as means ± S.E.M, n = 9-10 flies (except for Akh^-/-^ ND 3 weeks, where we used 20 flies) for each treatment from three replicates (***p < 0.001, as assessed by one way ANOVA followed by Tukey’s multiple comparisons).

## 4. Discussion

In this work, we recorded physiological and behavioral effects of short-as well as long-term exposure to HFD in *Drosophila.* We demonstrate that both virgin and mated female flies exposed to HFD exhibit decreased lifespan, and accelerated decline of climbing ability with age over three to four weeks. Some effects of HFD were seen only for either virgins or mated flies (see Table 1). Thus, an increase of sleep fragmentation after three weeks of HFD (30%) was shown only in virgin flies (mated ones were not tested; see section 3.2). Another long-term effect of HFD seen in virgin flies is a decline in TH immunolabeling in dopaminergic brain neurons. We detected no short or longterm effects of HFD on levels glycogen or glucose in flies of either condition, but in the short term (seven days) body weight decreased and TAG increased in mated flies. An important finding was that exposure to HFD for one or three weeks reduced egg laying and decreased egg numbers in the ovaries of mated flies. Furthermore, exposure to HFD for three weeks or longer lead to increased size of the crop and an elevation of *Akh* transcript levels. Since *Akh* transcript was upregulated after five weeks of HFD we analyzed the role of AKH signaling in longevity and fecundity under HFD conditions. *Akh* mutant flies displayed increased lifespan compared to controls both on ND and HFD. These mutants had reduced numbers of mature eggs in ovaries after one week on ND compared to controls, and a larger number after three weeks, but HFD decreased the egg number in both mutants and controls.

To the best of our knowledge, the novel findings in our study are the effects of HFD on crop size, sleep quality in ageing flies, fecundity, AKH signaling and dopamine signaling. Effects of obesity on *Drosophila* fecundity were analyzed earlier using obesogenic diets, not HFD, or by genetic interventions [see (Musselman and Kühnlein, 2018)]. Previous studies have also shown that HFD decreases lifespan and climbing ability (Jung et al., 2018; Rivera et al., 2019; Woodcock et al., 2015). Deleterious effects of HFD on several other behaviors during ageing have been reported, including memory and odor sensitivity (Jung et al., 2018; Rivera et al., 2019). Similar to humans, aging flies display decreased activity rhythms and increased sleep fragmentation (Koh et al., 2006; Metaxakis et al., 2014; Umezaki et al., 2012). In this study, we found that during nighttime, aging flies exhibited an increased number of sleep bouts, suggesting increased sleep fragmentation. We also noted a decrease in total duration of nighttime sleep. Additionally, we explored the effects of HFD and found that 30% HFD increased both total sleep amount and total number of sleep bouts in three-week-old flies compared to flies on ND. Thus, flies kept three weeks on 30% HFD sleep more than ND flies, but display increased sleep fragmentation. Interestingly, we also found that flies kept for three weeks on 30% HFD displayed hyperactivity during the first night after transfer to the DAMS system. This observation is consistent with a recent study (Huang et al., 2020), in which the authors showed that HFD increased starvation-induced hyperactivity. In conclusion, long-term exposure to HFD has an adverse effect on sleep.

Earlier studies found that short-term HFD induces increased TAG levels in flies (Birse et al., 2010; Diop et al., 2017; Jung et al., 2018). Our data show that virgin females kept on HFD for up to three weeks display no significant changes in body weight, TAG, glycogen or glucose levels. However, in mated females kept on HFD for one week there was an increase in TAG and over the first three weeks their body mass was lower than flies on ND. This diminished body mass is likely to depend on decreased oogenesis since we found lower numbers of eggs in mated flies on HFD.

In flies kept on 30% HFD for three weeks, the crop is drastically enlarged and filled with lipid. The crop is used for temporal storing of food, especially carbohydrates (Stoffolano and Haselton, 2013). Thus, it is important for maintaining carbohydrate homeostasis in flies. Normally the crop distension is sensed by proprioceptors that signal to brain circuits that regulate food intake (Wang et al., 2020), and there is also a neuronal and hormonal control of crop emptying into the gut (Gough et al., 2017; Kuraishi et al., 2015; Solari et al., 2017; Stoffolano and Haselton, 2013). In our experiments, it seems that HFD causes a defect in crop emptying leading to excess storage of ingested food, especially lipids. Coconut oil consists mainly of saturated fat in the form of free fatty acids (FFAs) like lauric or myristic acid (Birse et al., 2010). The FFAs seem to get trapped in the crop, and thus do not get converted to TAG. This could partly explain why long-term intake of HFD does not affect TAG levels in our assay system, which monitors oxidized glycerol after enzymatic hydrolysis of TAGs and does not detect FFAs. As mentioned in the introduction, it is important to note that the natural food of *Drosophila* is low in TAG and FFAs. Instead, a major dietary source of stored lipids is carbohydrates, which are converted into TAG in the fat body and intestine (Heier and Kühnlein, 2018). Thus, the crop may fail to efficiently redistribute excess FFAs. The crop malfunction may also affect ingestion of other nutrients and alter metabolism. However, only in the mated flies we noted an effect of HFD on body mass and TAG levels, whereas glycogen and glucose levels were not affected. In virgin flies there was no significant effect on any of these parameters. Possibly alterations in nutrient uptake and metabolism lead to reallocation of energy for somatic maintenance in females (Partridge et al., 1987). This could explain the decreased fecundity in mated flies kept on HFD (seen both as a decrease in mature eggs in the ovaries and reduced egg laying).

Long-term exposure to HFD resulted in increased *Akh* transcript levels, although no change was seen after short exposure, and the AKH peptide levels remain unaltered. These findings may suggest that in the long-term AKH production and release increases in response to HFD. Another study revealed increased AKH receptor transcript after exposure to HFD (Huang et al., 2020) and taken together with our data, AKH signaling thus appears to increase on this diet. AKH is known to induce mobilization of lipids in insects, including *Drosophila* (Gäde and Auerswald, 2003; Grönke et al., 2007; Musselman and Kühnlein, 2018; Toprak et al., 2020; Van der Horst, 2003), but also to allocate carbohydrate as energy to fuel food search in hungry flies (Bharucha et al., 2008; Isabel et al., 2005; Lee and Park, 2004; Yu et al., 2016). AKH signaling has been shown to regulate lifespan in flies (Katewa et al., 2012; Post et al., 2019; Waterson et al., 2014). More specifically, ubiquitous overexpression of *Akh* (Katewa et al., 2012), long-term thermogenetic activation of AKH producing cells and mutation of the water sensing protein pickpocket 28 (*ppk28*) (Waterson et al., 2014) or mutation of *dilp2* (Post et al., 2019) lead to increased *Akh* levels and a prolongation of life span. In our work, we analyzed female *Akh^AP^* mutant flies (also lacking the AKH precursor related peptide, APRP) and found that both on ND and HFD the mutants lived longer than *w^1118^* controls, although HFD shortened lifespan both for mutants and controls. A previous analysis of *Akh* mutant flies (only defect in *Akh*) detected no significant effect on lifespan in females or males (Bednářová et al., 2018). Thus, there are conflicting results from experiments with *Akh* mutants and *Akh* upregulation on lifespan in these studies. Possibly some of the differences can be attributed to variations in experimental protocol and diets. In this context, it can be noted that *Akh* mutant flies live longer when exposed to starvation (Isabel et al., 2005).

AKH is required for mobilization of energy substrates, notably lipid and trehalose, during energy-demanding exercise, such as flight and locomotion, including starvation-induced food seeking (Gäde and Auerswald, 2003; Isabel et al., 2005; Lee and Park, 2004; Van der Horst, 2003; Yu et al., 2016). *Akh* expression increased almost four-fold after five weeks of HFD compared to ND and short-term exposure to HFD. This may suggest that five weeks of HFD renders flies that experience nutrient deficiency. In fact, when flies are transferred from three weeks of HFD to activity monitors, they display increased activity the first 12 hours (**Fig. 3B**), which may represent hunger-mediated food seeking behavior (subsequently the flies feed on sucrose-yeast diet in the monitor tubes). We also showed that flies kept on HFD for a week and then transferred to the CAFE assay ingested significantly more food the first 24 hours than controls on ND. Taken together, the increased AKH signaling, increased locomotor activity and food intake, suggest that flies on HFD experience an energy shortage.

Although several of the DILPs are involved in lipid metabolism in flies (Broughton et al., 2005; Musselman and Kühnlein, 2018; Toprak et al., 2020), we found that transcript levels of *dilp1-dilp3, dilp5,* and *dilp6* displayed no significant changes with short of long-term exposure to HFD. Similar findings were presented in previous reports (Jung et al., 2018; Woodcock et al., 2015). However, we observed a significant decrease in DILP2 peptide levels in the IPCs after one week of HFD, but not after five weeks, suggesting short-term effects to trigger DILP2 release. DILP2 is one of the peptides released from IPCs upon food intake, and it was recently shown that the brain IPCs are triggered by activation of the Piezo proprioceptors of these cells upon distension of the crop (Wang et al., 2020).

What are the mechanisms behind the shortened lifespan and behavioral senescence in flies kept on HFD? An excess of dietary lipids affects several tissues such as the intestine, the fat body, the central nervous system, and the heart, but also results in global effects on metabolism and the expression of a multitude of genes (Birse et al., 2010; Huang et al., 2020; Jung et al., 2018; Musselman and Kühnlein, 2018; Rivera et al., 2019; von Frieling et al., 2020; Woodcock et al., 2015). Some effects of HFD seem to be tissue specific, others are more global.

The heart function in *Drosophila* deteriorates already after short exposure to HFD with the appearance of lipid accumulation, reduced contractility and structural changes in heart muscle (Birse et al., 2010). It therefore seems that, at least in the short term, HFD has direct lipotoxic effects on the heart muscle. These HFD effects can be alleviated by both local and systemic reduction of activity in the insulin-TOR (target of rapamycin) pathway (Birse et al., 2010). This suggests that HFD causes a deregulation of the insulin-TOR pathway, at least in the short-term. Surprisingly, we noted no anatomical long-term effects of HFD on the heart. At the gut level, HFD alters the microbiota, increases intestinal stem cell activity, and increases the number of enteroendocrine cells in the midgut (von Frieling et al., 2020). In that study, it was shown that the HFD directly activates JNK signaling in enterocytes that in turn signal with the cytokine unpaired 3 (upd3) to activate JAK/STAT signaling in the stem cells. Especially in the long-term this signaling is dependent on presence of microbiota. The HFD induced changes in the gut function are likely to alter both intestinal and organismal metabolic homeostasis (due to peptide hormones in enteroendocrine cells) and also ageing of the gut. Another study implicated HFD-activated upd3 and JAK/STAT signaling in macrophage activation and inflammation in the shortened lifespan and altered metabolism of *Drosophila* (Woodcock et al., 2015). Other studies have suggested that HFD alters mitochondrial respiration in cells, including neurons, that may result in behavioral senescence (Cormier et al., 2019), or induces effects on specific neurons via the lipoprotein LPT and its receptor LpR1 to induce behavioral changes (Huang et al., 2020). Two studies investigated HFD-induced alterations in gene expression that affected olfactory behavior, metabolism, and stress responses (Jung et al., 2018; Rivera et al., 2019). It is also clear that obesity, caused by factors other than HFD, influences ovary development and fecundity (Musselman and Kühnlein, 2018). Thus, in summary, there are several mechanisms that can explain the effects of HFD on longevity, fecundity, and behavioral senescence.

## 5. Summary and conclusions

In summary, our data suggest that exposure to HFD not only reduces lifespan, and accelerates age-related deterioration of behaviors such as negative geotaxis and sleep patterns, it also leads to reduced fecundity. Furthermore, HFD leads to increased food intake when transferred from HFD to ND, and induces short-term hyperactivity that indicates hunger-driven food seeking; taken together these two findings suggest that flies experience an energy deficiency. Mated females, but not virgins, exposed to HFD display decreased body weight and increased TAG levels compared to controls. This weight difference can probably be explained by the reduced number of mature eggs in the ovaries of HFD flies. Furthermore, *Akh* expression, and probably AKH signaling, is elevated in flies after long-term HFD suggesting increased energy mobilization. Lifespan is extended in *Akh* mutant flies kept on HFD compared to control flies on the same diet, but it is shortened compared to mutants kept on ND. Thus, taken together our findings indicate that AKH signaling plays a role in the response to HFD. The enlarged, lipid-filled crops observed in flies on HFD, are remarkable and suggest a malfunction in nutrient allocation in the fly that may contribute to the HFD morbidity.

## Acknowledgements

We thank Dr. Susan Broughton for advice early in this project and Drs John Stoffolano and Alan Thomson for stimulating discussions on crop function. We thank the following people and organization for flies and reagents: Bloomington *Drosophila* Stock Center (BDSC), Bloomington, IN, USA (supported by NIH P40OD018537), Drs Ronald Kühnlein, Mark R. Brown and Jan A. Veenstra (see Material and methods for details). We are grateful to the Swedish Research council (Vetenskapsrådet) and Carl Trygger Foundation for funding. Stina Höglund and the Imaging Facility at Stockholm University (IFSU) are acknowledged for maintenance of the confocal microscopes.

## Supplementary material figures

**Supplementary figure 1.** Experimental design. **A.** Regime 1: Virgin female flies were collected after eclosion, and were thereafter kept either on normal diet (ND) or food containing 10% coconut oil (HFD 10%) or food containing 30% coconut oil (HFD 30%). **B.** Regime 2: Flies were allowed to mate on ND for 4 days after eclosion, thereafter these flies were transferred to ND or HFD 30%. The time line is shown as a horizontal bar, where Ecl depicts collection of newly eclosed flies and 0 the transfer to experimental diet. The effects of HFD on lifespan of virgin and mated females were determined, and assays were performed at the sample times indicated.

**Supplementary figure 2.** Effects of HFD on climbing and chill coma recovery. **A.** Test of negative geotaxis (climbing assay). Mated flies kept for 3 to 4 weeks on HFD displayed decreased climbing ability compared to ND controls. Data are presented as means ± S.E.M, n = 6 replicates with 10 flies in each replicate for each group (*p < 0.05, two-way ANOVA followed by Sidak’s multiple comparisons test). **B.** Chill coma recovery in flies on HFD. Virgin flies kept for one week on 30% HFD recovered more rapidly from chill coma compared to ND controls. Data are presented as means ± S.E.M., 80-94 flies per treatment divided in three biological replicates were used group (***p < 0.001, one-way ANOVA followed by Tukey’s multiple comparisons).

**Supplementary figure 3**. The average locomotor activity over 3 days of virgin female flies fed with ND and HFD. Flies fed with 10% HFD display no change in locomotor activity after 1 week (**A**) nor 3 weeks (**C**) on the diet. However, flies fed 30% HFD decreased their locomotor activity both after 1 (**B**) and 3 weeks (**D**) on the diet, especially during the light phase (**D**). **E.** Summary of data depicted in A-D shown as total activity/day (average activity per 24-hour period across three days). The 30% HFD flies decreased total activity compared to ND flies at both 1 and 3 weeks of treatment. Data are presented as means ± S.E.M, n = 46-48 flies for each treatment from three replicates (*p < 0.05, as assessed by one-way ANOVA followed by Tukey’s multiple comparisons).

**Supplementary figure 4**. Effects of HFD on the aging of dopamine neurons and muscle in virgin flies. **A.** Representative images of TH immunolabeling in dopaminergic protocerebral posterior lateral (PPL1) neurons after 1 week ND, 3 weeks ND and 3 weeks HFD. **B.** Tyrosine hydroxylase (TH) immunolabeling diminished in PPL1 dopamine neurons in flies kept 3 weeks on HFD compared to 1 and 3 weeks on ND. Data are presented as means ± S.E.M, n = 8-10 flies for each treatment from three replicates (***p < 0.001, as assessed by one-way ANOVA followed by Tukey’s multiple comparisons). **C.** Images depicting polyubiquitin immunolabeling in abdominal muscles of flies fed ND and 30% HFD for 1 and 5 weeks. **D-F.** Quantification of polyubiquitin immunolabeling. The number, average size and the percentage area of polyubiquitin stained particles were quantified. HFD does not affect the accumulation of polyubiquitin immunolabeling, only age. Data are presented as means ± S.E.M, n = 10-11 flies for each treatment from three replicates (*p < 0.05, **p < 0.01, ***p < 0.001, as assessed by one-way ANOVA followed by Tukey’s multiple comparisons). **G.** The morphology of the abdominal heart muscle (phalloidin staining) is affected by age over five weeks, but not by 30% HFD.

**Supplementary figure 5**. Adipokinetic hormone (AKH) mutant phenotypes. **A** and **B**. AKH immunolabeling is totally abolished in the *Akh* mutants (Akh^-/-^) compared to their controls (w^1118^). **C.** Crops of some *Akh* mutants (mated flies) increase in size even on normal diet (ND). These crops do not contain Nile Red stained lipid aggregates, in contrast to enlarged crops from mutants kept on high fat diet (compare with Fig. 8D).

