## Supplementary figures and images for "Impact of high-fat diet on lifespan, metabolism, fecundity and behavioral senescence in *Drosophila*"

### Supplemental Figure 1

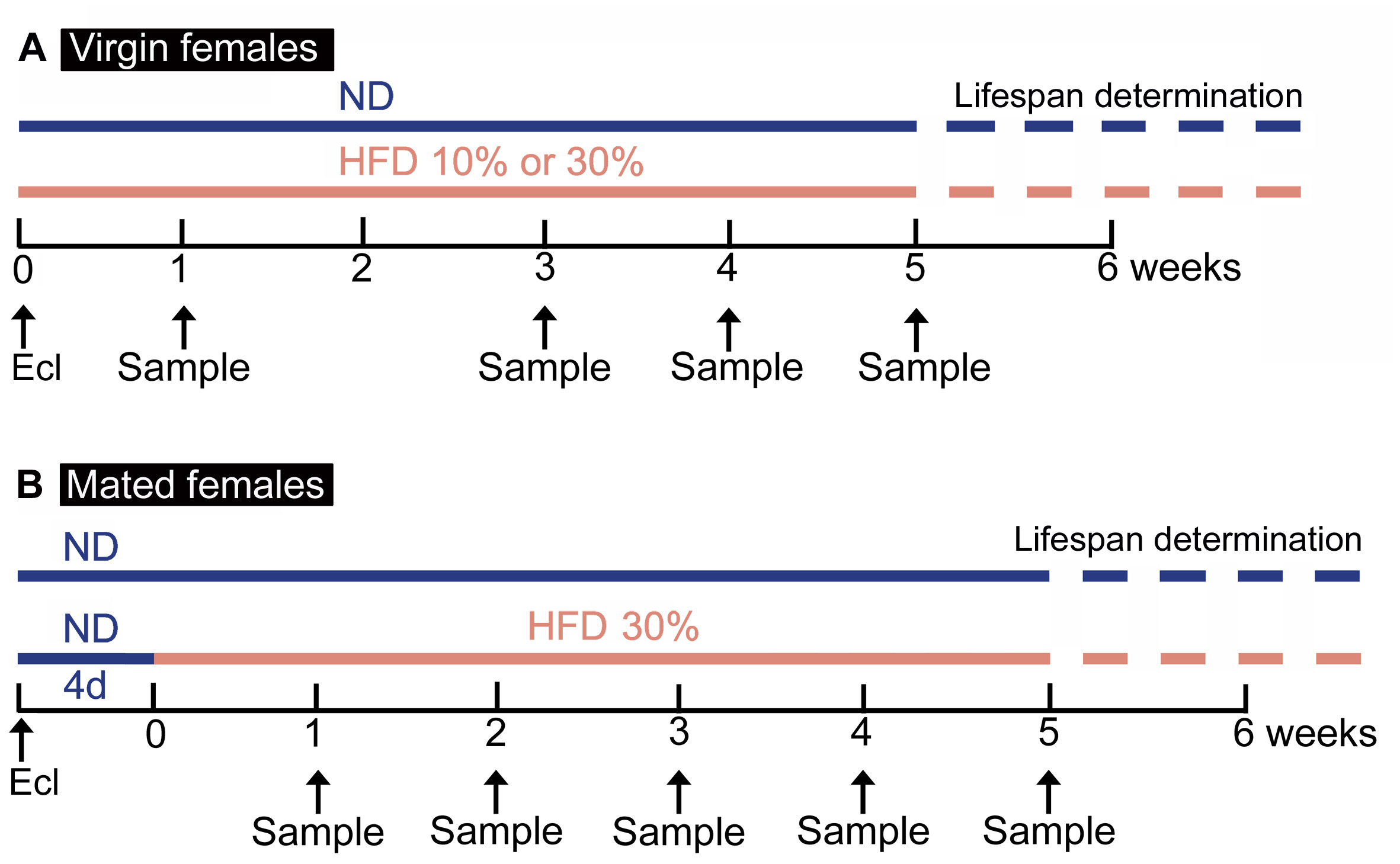

### Supplemental Figure 2

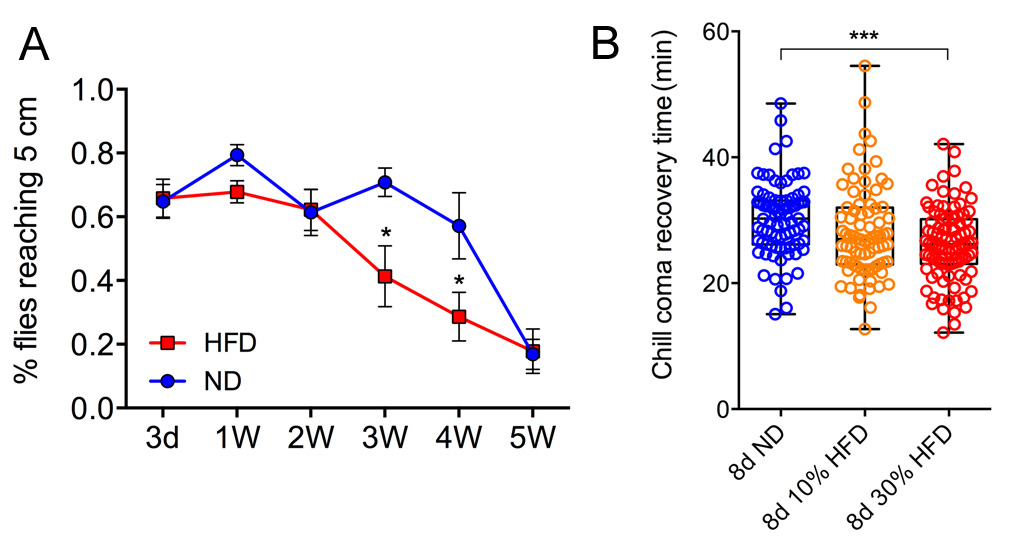

### Supplemental Figure 3

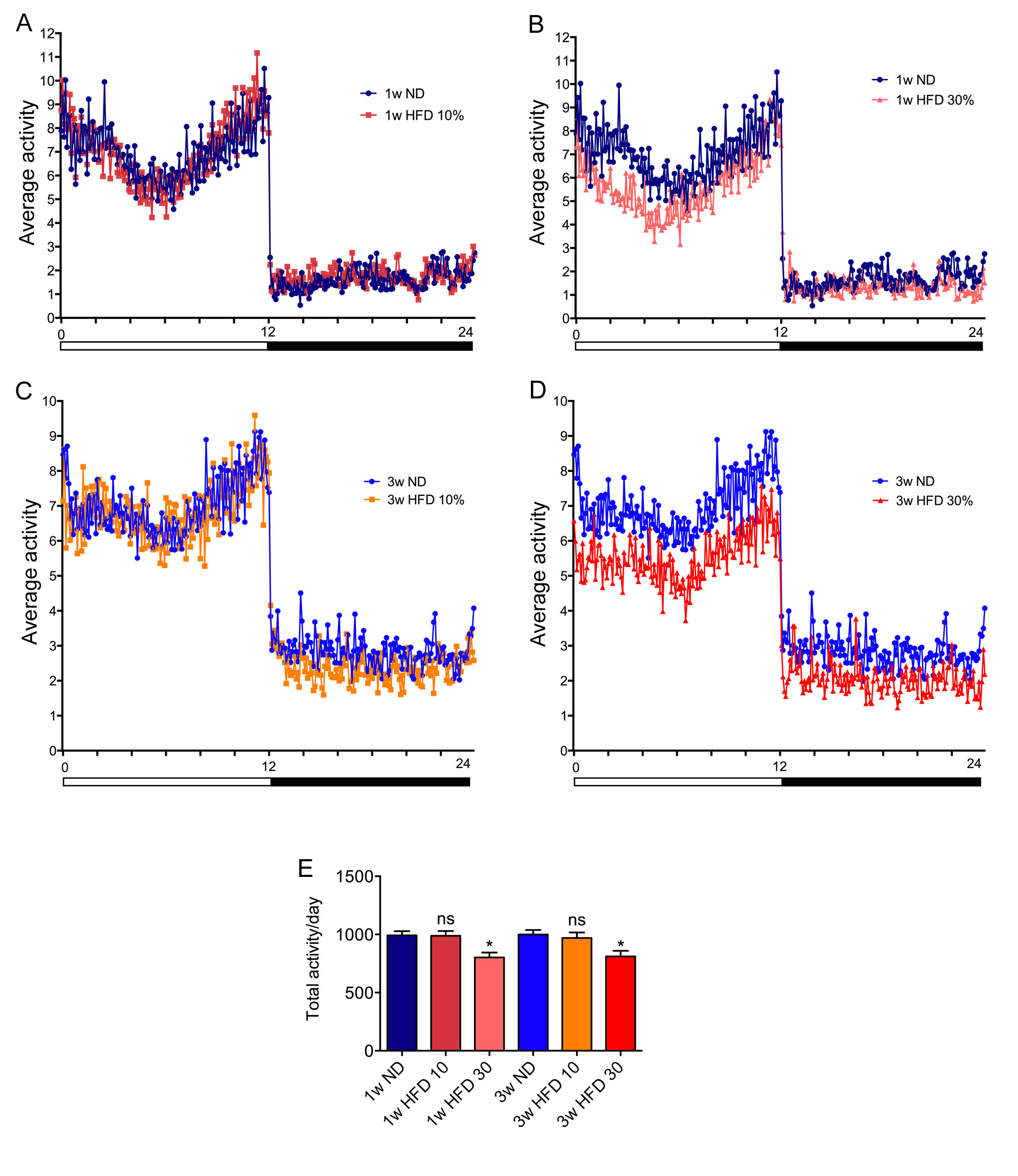

### Supplemental Figure 4

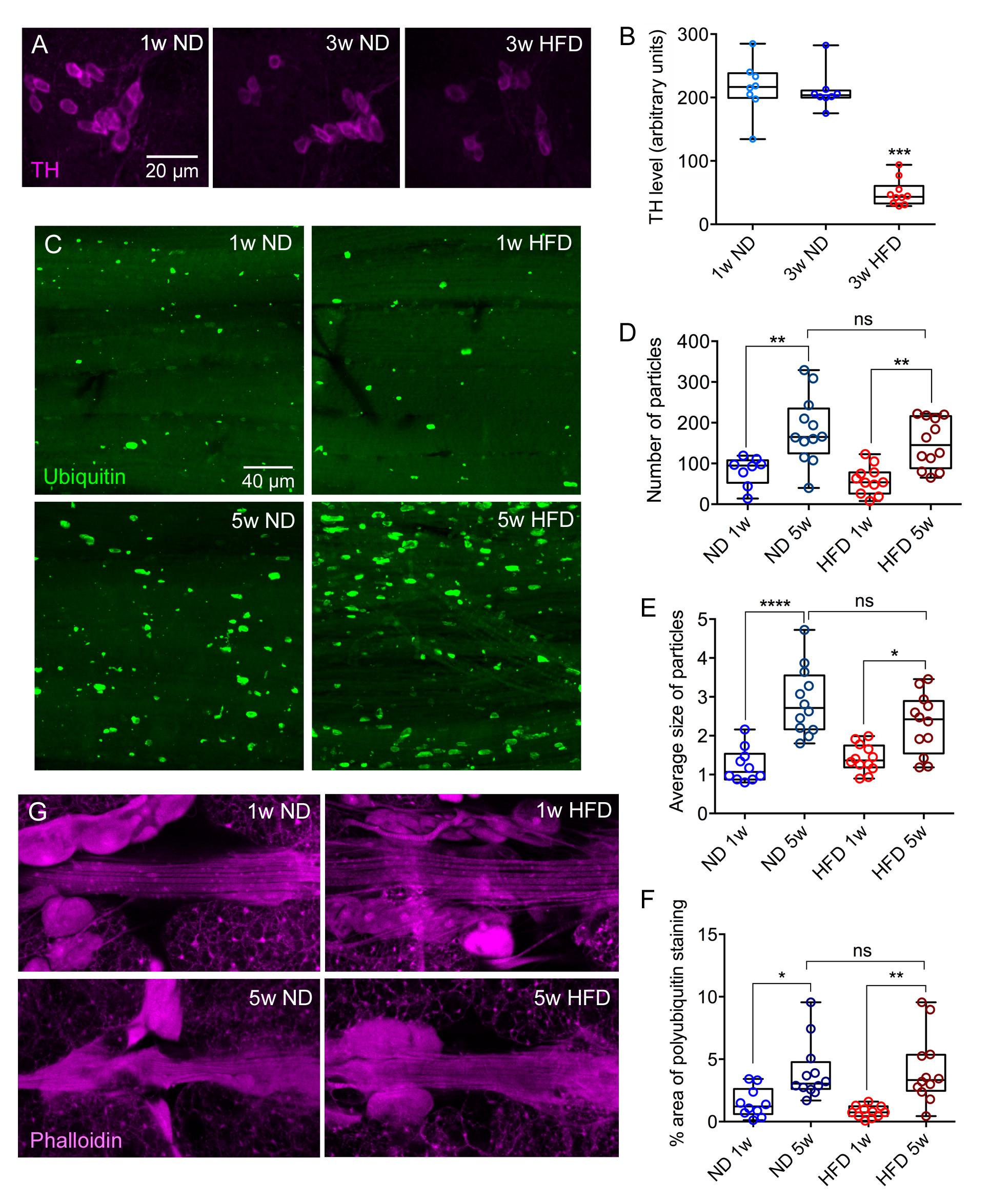

### Supplemental Figure 5

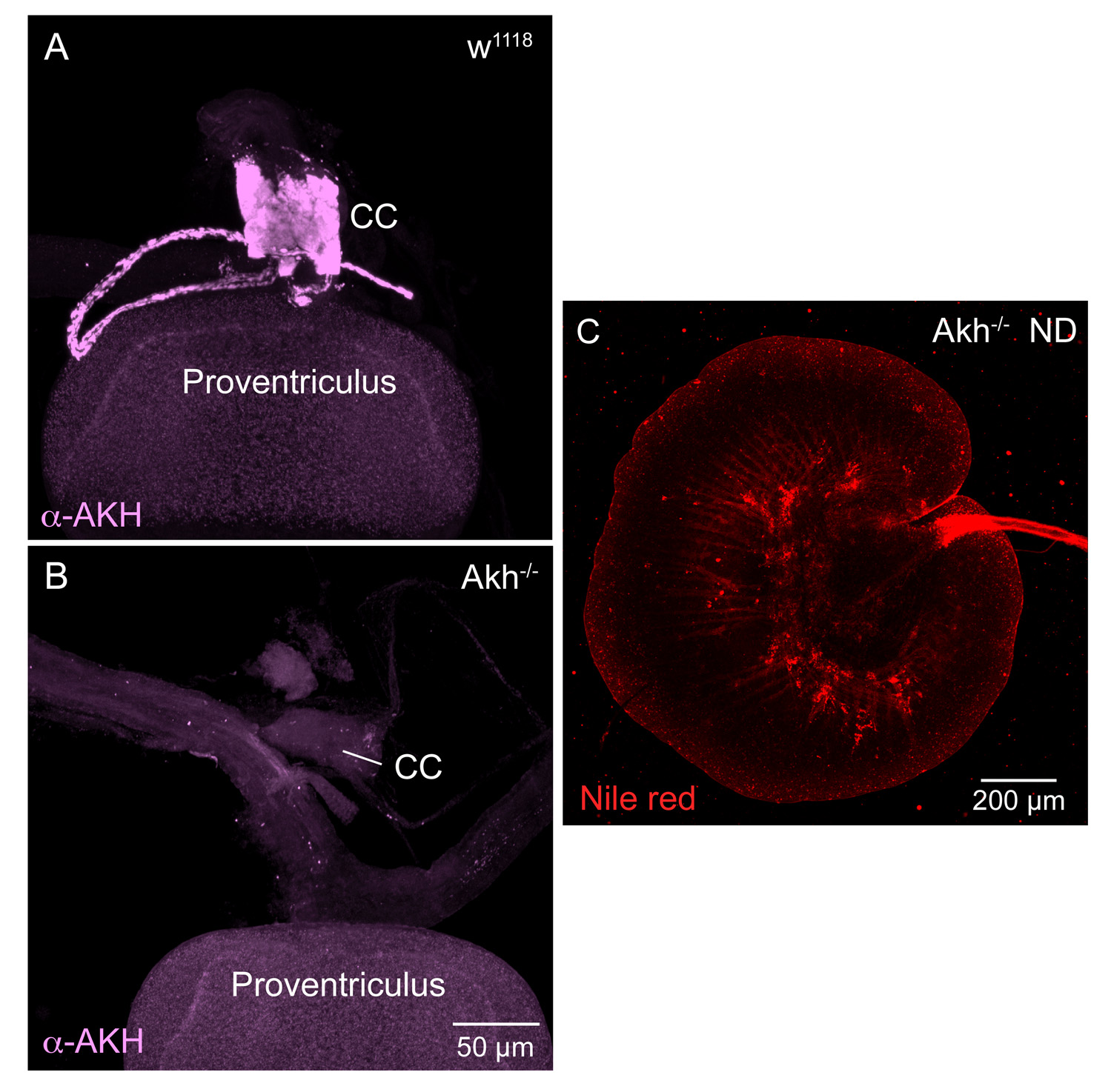
